# Age and Sex-Dependent Differences in Human Cardiac Matrix-Bound Exosomes Modulate Fibrosis through Synergistic miRNA Effects

**DOI:** 10.1101/2022.11.14.516464

**Authors:** George Ronan, Gokhan Bahcecioglu, Jun Yang, Pinar Zorlutuna

## Abstract

Aging is a risk factor for cardiovascular disease, the leading cause of death worldwide. Cardiac fibrosis is a harmful result of repeated myocardial infarction that increases risk of morbidity and future injury. Interestingly, rates of cardiac fibrosis are different between young and aged individuals, as well as men and women. Here, for the first time, we identify and isolate matrix-bound extracellular vesicles from the left ventricles (LVs) of young or aged men and women. These LV vesicles (LVVs) show differences in morphology and content between these four cohorts. LVVs effects on fibrosis were also investigated *in vitro*, and it was shown that aged male LVVs were pro-fibrotic, while other LVVs were anti-fibrotic. miRNAs identified from these LVVs could partially recapitulate these effects together, but not individually, and confer other benefits. These data suggest that synergistic effects of matrix-resident exosomal miRNAs may influence the differential clinical response to MI.

## INTRODUCTION

Cardiovascular disease (CVD) is the leading cause of death in the United States and worldwide, with myocardial infarction (MI) as the chief cause of death among CVDs^1^. While initial incidence of MI tends to be non-fatal, the tissue response results in thus-far irreversible damage to the myocardium^2^. This damage commonly takes the form of cardiac fibrosis, or excessive scarring and defunctionalization of the cardiac tissue, which increases risk and mortality of a future cardiac event^2, 3^.

Aging is a major risk factor for cardiovascular disease and numerous other diseases and is a growing area of research given the aging population in the United States and other countries^4, 5^. Furthermore, data increasingly suggests that age and sex play significant roles in the likelihood and severity of MI and resulting fibrosis^5–8^. Males over 50 years of age tend to have a higher risk of fibrosis and typically experience MI 9 years earlier than females, although these differences subside as age surpasses 80 years^6, 9, 10^. While the precise reasons for these discrepancies remain elusive, recent data suggests that aging and sex-related long-term changes to the cardiac microenvironment account for this differential response to MI^6, 8, 11^.

In recent years, the use of extracellular matrix (ECM) or ECM-derived materials in the treatment of cardiac fibrosis has seen reliable success in pre-clinical trials^12–16^. These approaches take advantage of the endogenous cardioprotective effects of ECM on the local microenvironment to promote functional tissue recovery after cardiac injury^17, 18^ through local immunomodulation, stem cell recruitment, and decreased scar tissue formation^19–21^. These effects synergize to enhance regenerative healing and decrease fibrosis post-MI. However, the mechanisms by which ECM promotes cardiac repair are not well understood, and recent studies suggest that the release of embedded signaling molecules such as cytokines or growth factors^22^ and ECM-microRNA (miRNA) interactions^11^ are major effectors of both pro and anti-fibrotic signaling pathways post-MI.

The identification of these factors suggest that the beneficial effects of ECM may be conferred by extracellular vesicles (EVs), as both cytokines and miRNAs are commonly packaged in EVs when secreted from cells^23^. Previously unidentified ECM-bound EVs could be key mediators of the beneficial effects of ECM treatment, and isolation, quantification, and characterization of these EVs will elucidate essential mechanisms of ECM-mediated cardioprotection. Furthermore, the isolation of key functional compounds, either EVs themselves or those contained in EVs, may provide the benefits of ECM treatment while mitigating many associated challenges, such as immune response, sample preparation variability, and sustainability of production. Another challenge, however, is how to modulate key regulators of fibrosis-related signaling pathways after identifying them. While cytokine-mediated inflammatory signaling pathways are attractive targets for clinical intervention, as they play a pivotal role in the health and functionality of a tissue and allow for direct intervention in the onset and resolution of inflammation^24, 25^, the degree and mechanisms of involvement remain an active area of research^24^. Recent advances in our understanding of the tissue microenvironment *in vivo* have suggested that this may be due to targeted paracrine signaling controlling these effects, resulting from the highly specific packaging of miRNAs and cytokines in exosomes, a specific, tightly regulated class of EV^23^.

Exosomes are a subgroup of EVs with diameters typically between 30 nm and 200 nm that are commonly released from most cell types and contain cytokines, chemokines, miRNA, and other miscellaneous signaling molecules that affect function in recipient cells. These contents influence many diverse and pathologically relevant biological processes, including angiogenesis, immunomodulation, endothelial and epithelial to mesenchymal transition, and cell differentiation, and as such exosomes are both packaged and released from cells in a highly controlled manner^23, 26, 27^. As a result, recent interest in exosomes has primarily been in the role of maintaining tissue health through intra-tissue signaling and local immunomodulation^23, 27^. This has been bolstered by the recent discovery of exosome-like EVs embedded in decellularized tissue from several human organs, including the urinary bladder and small intestine^26^, and decellularized mouse atrium^28^, as opposed to biofluid-derived EVs which have so far been ubiquitous. These embedded EVs showed beneficial immunomodulatory effects, and those isolated from cardiac tissue enhanced cardiomyocyte function *in vitro*^28, 29^, which provides exciting prospects for how these EVs may affect MI response and subsequent cardiac fibrosis and suggests that these EVs may be a core functional component of biosignaling in the ECM. Further investigation of these EVs may reveal precise mechanisms by which ECM treatment confers protective effects, both furthering knowledge of the interplay between microenvironment and tissue health and providing a wealth of targets for clinical intervention without necessitating the use of ECM. However, the impact of both induced and innate differences in the microenvironment may have on these ECM-bound exosomes is currently unknown.

Recently, exosomes have been increasingly investigated for links with MI response^14^. Exosomes isolated from the cerebrospinal fluid or plasma of young or aged subjects have been demonstrated to have differential effects on modulation of systemic inflammation and progression of neurodegenerative diseases^4, 30^, and these differences can significantly affect CVD outcomes *in vivo*^30^. However, despite evidence suggesting that there are functional differences between exosomes from young or aged subjects, there has been little evaluation of the specific differences between these exosome populations. For this reason, exosomes have become an attractive target for ascertaining specific age-related changes in the cardiac microenvironment and the impact of any additional factors.

In this study we show for the first time in literature the presence of ECM-bound exosome-like EVs in human left ventricular (LV) tissue, and report the changes in size, cytokine content, and miRNA content of these LV vesicles (LVVs) as a function of age and sex. Furthermore, we study the differential effect of LVVs derived from different age and sex groups on the stress response and fibrotic transdifferentiation of cardiac fibroblasts (CFs) to myofibroblasts (MFs) in both human and murine models. Following this, we examine the role of select miRNAs identified from the LVVs in modulating this transdifferentiation. In this way, we suggest that ECM-bound exosomes are a major functional unit of the cardioprotective effects of the ECM, hosting previously identified signaling molecules of interest and recapitulating the effects of ECM treatment on the local microenvironment. Investigating the effects of age and sex on the physical characteristics and composition of human LVVs and how LVVs influence the fibroblast transdifferentiation behavior as a function of age and sex will pave the way for understanding the mechanisms of cardiac fibrosis and developing new treatment strategies to prevent fibrosis and MI.

## RESULTS

### Left Ventricular Vesicle (LVV) Isolation and Characterization

Human heart left ventricular tissues from young female (YF), young male (YM), aged female (AF), or aged male (AM) donors were subjected to a detergent-free decellularization and EV isolation process (Figure 1A). Transmission electron microscopy (TEM) imaging verified the presence of EVs in the ECM isolate and demonstrated a stark size difference between young and aged tissue-derived vesicles (Figure 1B). Similar properties were observed in tissue examine directly under TEM (Supplemental Figure S1). This difference was also observed with nanoparticle tracking analysis (NTA), with aged EVs having average size of 98 nm ± 22 nm and young EVs having average size of 171 nm ± 38 nm (Figure 1C). All samples fell primarily within the expected size range for exosomes (30-200 nm), and dispersity decreased in aged tissue-derived samples compared to young. The concentration of vesicles was not significantly different between cohorts (Figure 1D). Western blot was performed to identify characteristic exosome markers CD9, CD63, and TSG101 (Figure 1E), as well as for characteristic cardiac cell markers Vimentin, CX43, and VE-Caherin (Supplemental Figure S2). These results showed that the particles were, or contained, exosomes, and were free of cell debris. Lysing these vesicles increased the measured protein content in solution by over 50% (Figure 1F), indicating that the isolated LVVs contained proteins.

**Figure 1.**
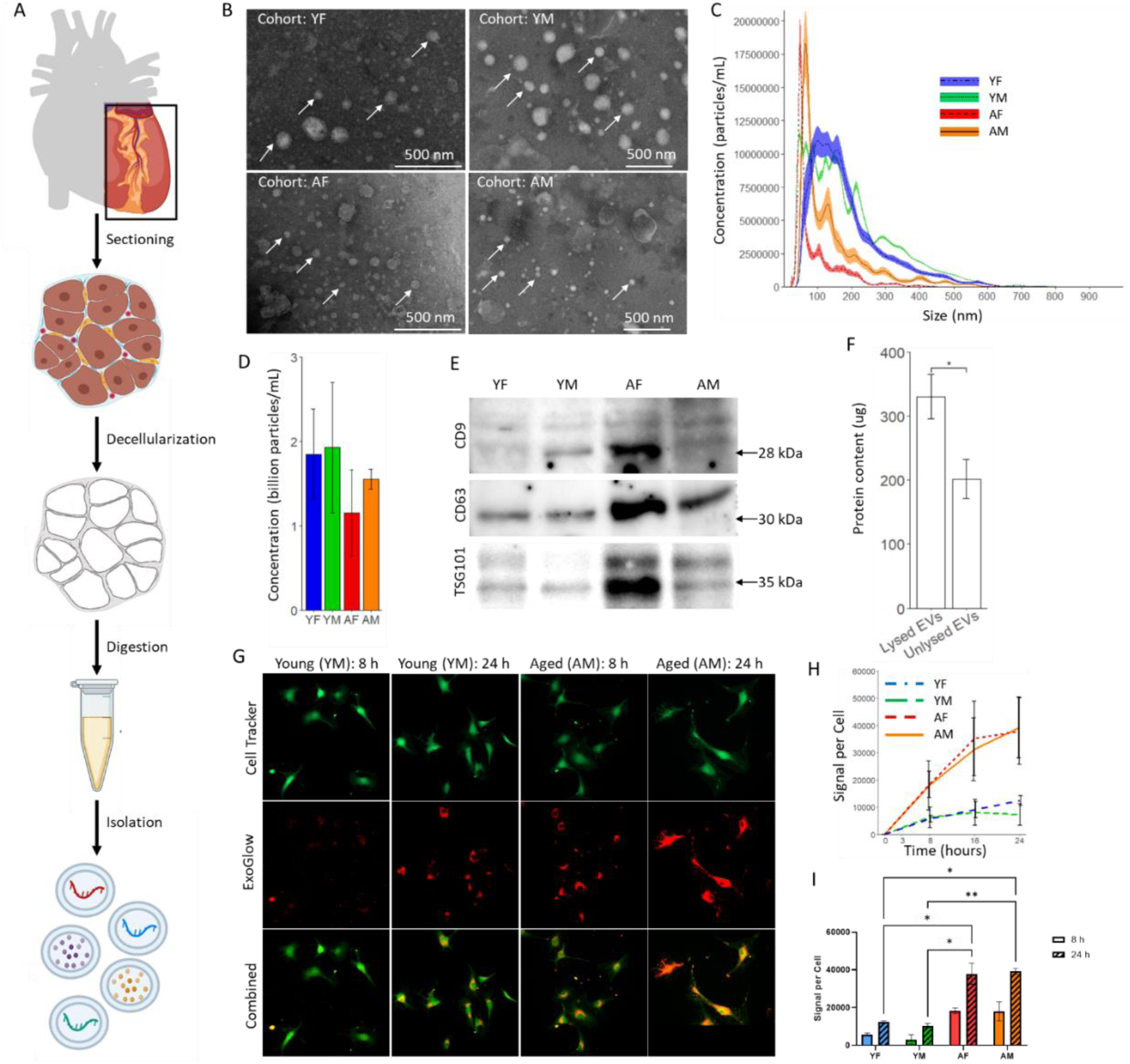
Functional Exosomes can be Obtained from Human Left Ventricular Extracellular Matrix with Distinct Aging-Related Changes. (A) Brief overview of the ECM-bound EV isolation process. (B) Transmission electron microscopy for representative imaging of all four cohorts. (C) Nanoparticle tracking analysis with error area of EVs from all four cohorts. (D) Typical concentration of LVVs per sample. (E) Western blotting of lysed LVVs showing the characteristic exosome markers. (F) Protein encapsulation within LVVs. (G) Representative images of stained EV uptake by hCFs, with corresponding (H) uptake rate and (I) overall uptake by cells (n ≥ 3 for all LVV sources in all cohorts, 5 images per sample). Data are presented as the mean ± standard deviation. * p < 0.05, ** p < 0.01, assessed by Student’s t-test with Welch’s correction for (D), (F), and (I).

### LVV Uptake by Human Cardiac Fibroblasts (hCFs)

Uptake of EVs by hCFs was confirmed by tracking stained EVs in cells (Figure 1G, Supplemental Figure S3). Aged EVs were taken up by hCFs at a nearly 2-fold increased rate compared to young EVs (Figure 1H), with a nearly 2-fold increase in concentration of aged EVs per cell from 8 to 24 h post treatment, and only up to 1.5-fold increase in concentration of young EVs (Figure 1I). Higher overall quantities of aged EVs per cell were also taken up compared to young EVs, with aged EVs being taken up at 2 to 3-fold higher quantity than young EVs at all timepoints (Figure 1I).

### Effect of LVVs on 3D hCF Gel Contraction

To assess the effects of LVV treatment on MF transdifferentiation in 3D culture, hCFs were seeded either in collagen gels with gel contraction assessed, or on tissue culture plates with wound healing (through scratch assay) assessed (Figure 2A). For gel contraction assay, cells were evenly distributed throughout the gel during seeding and attached and spread within the gels (Supplemental Figure S4). All gels maintained structural stability for at least 48 h and no contraction was observed in the cell-free gel control (Figure 2B). The size of the cell-loaded gels from all groups followed a logarithmically decaying curve over time and, compared to the LVV-untreated control, the reduction in gel size was decreased after treatment with YF, AF, or YM LVVs and increased after treatment with AM LVVs (Figure 2C). The total contraction after 48 h was significantly higher in the AM LVV treated group, and significantly lower in the YF, AF, and YM LVV treated groups, compared to the untreated control (Figure 2D). The least contraction was observed with AF group.

**Figure 2.**
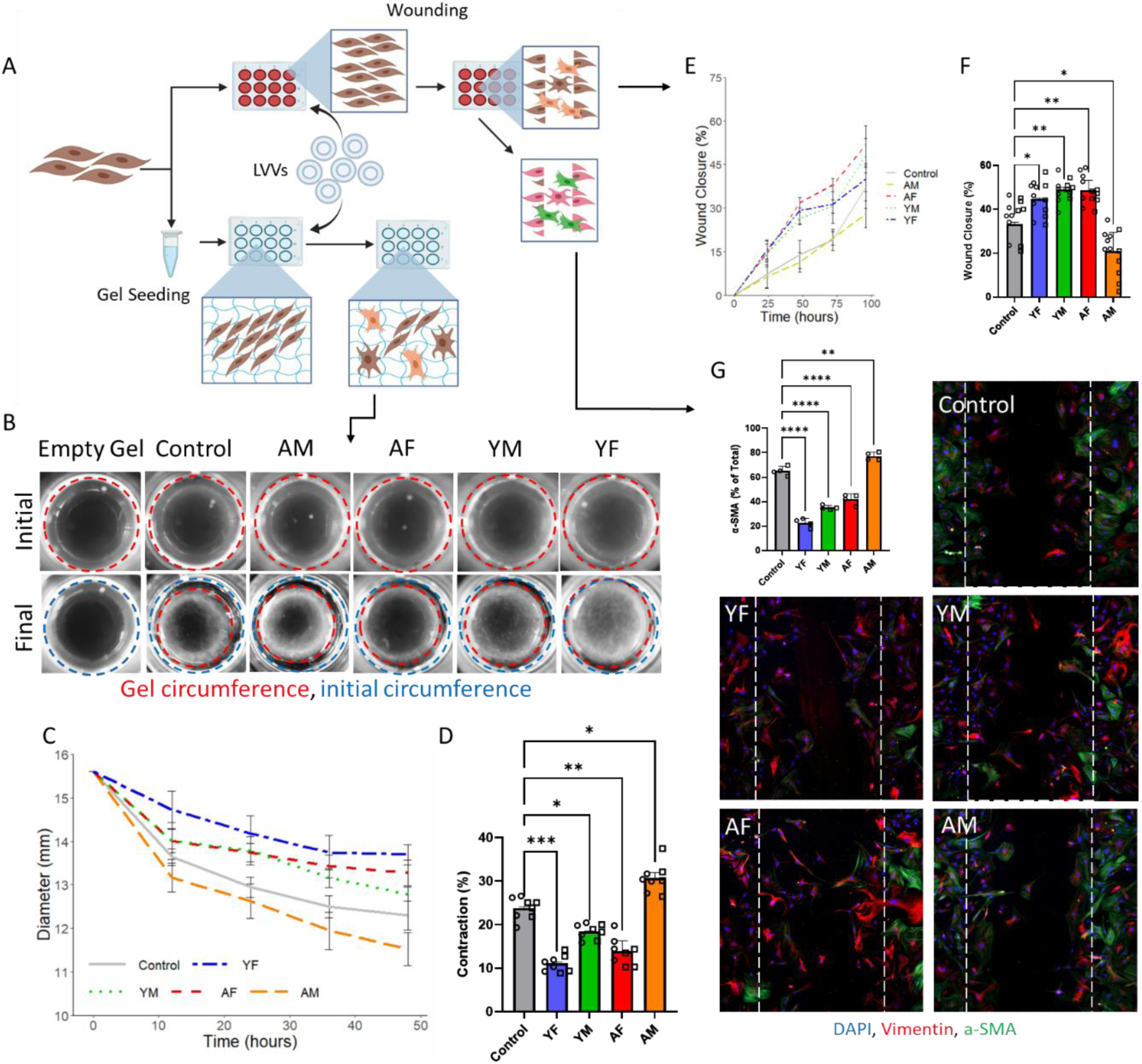
LVVs Modulate Fibroblast Behavior In Vitro to Control Transdifferentiation and Fibrotic Effects. (A) Schematic briefly showing the assays performed with LVV treatment and resulting data. (B) Representative images of gels for both baseline and final timepoints for each cohort with gel circumference indicated, quantified as rate of gel contraction over 48 h (C) and final contraction percentage calculated relative to initial gel diameter (D) for each cohort (n ≥ 3 for all LVV sources in all cohorts). Wound closure (E) rate and (F) percentage over 96 h for each cohort (n ≥ 3 for all LVV sources in all cohorts, 2 technical replicates per sample, 3 images per replicate, for 2 independent repetitions). (G) Percentage of cells expressing α-SMA for each cohort, and representative images of stained wound area (n ≥ 3 for all LVV sources in all cohorts, 3 images per sample). Data are presented as the mean ± standard deviation. Bar graphs represent biological replicates, with technical replicates overlaid as a dot plot. * p < 0.05, ** p < 0.01, *** p < 0.001, assessed by one-way ANOVA with Tukey’s post-hoc for (D), (F), and (G).

### LVVs Affect hCF Wound Healing and Transdifferentiation

Scratch assay demonstrated 1.5 or 2-fold enhanced wound closure over culture period after treatment of hCFs with YF, AF, and YM LVVs compared to the control, and 30% decreased wound closure after treatment with AM LVVs (Figure 2E). At the endpoint, all groups demonstrated significantly different wound closure behavior from the control, with AM LVVs having decreased closure while other groups having increased it (Figure 2F). A similar trend was observed with rat CFs (Supplemental Figure S5), although YF LVVs showed no beneficial effects on these cells. No group achieved full wound closure in the time allotted, with the YF, AF, and YM groups achieving >45% average closure, the control achieving ∼37% closure, and the AM group achieving <30% closure (Supplemental Figure S6). Immunostaining of cells at 96 h post treatment revealed significant differences in α-SMA expression between all groups and the control, with the YF, AF, and YM groups showing <40% α-SMA expression compared to ∼65% expression in the control and >75% expression in the AM group (Figure 2G). The differences observed inversely corresponded to wound healing capacity, with groups demonstrating enhanced wound healing capacity expressing lower levels of α-SMA and vice-versa (Figures 2F and 2G). Greater than 95% of cells in all groups expressed vimentin, confirming them as fibroblasts (Supplemental Figure S7).

### Profiling of Cytokine Content

Cytokine profiling via dot blot-based immunoassay revealed differential concentrations of cytokines present in LVVs from different subject groups. In general, when compared to YF LVVs, which had the measured lowest quantity of cytokines (Supplemental Figure S8), YM and AF LVVs showed little difference while AM LVVs showed over 3-fold higher levels of several cytokines (Figure 3A). These include Angiopoietin-2 (2.99-fold), Dkk-1 (2.58-fold), Emmprin (2.455-fold), IFN-γ (2.84-fold), IL-1α (3.05-fold), Kallikrein-3 (3.31-fold), and SDF-1α (2.94-fold) among others (Supplemental Table S1). Proteomapping of the affected KEGG pathways showed upregulation of transport and HIF-1, Ras, and TNF signaling, and downregulation of PPAR signaling in aged subjects compared to young, and in males compared to females (Figure 3B). Interestingly, cytosolic DNA sensing was observed in males but not females. Additionally, gene ontology analysis showed that cytokines present in the AM LVVs were associated with regulation of tissue remodeling, positive regulation of receptor-mediated endocytosis and cytokine production, and negative regulation of cell death and wound healing (Supplemental Table S2). Only male LVVs were involved in negative regulation of wound healing (Supplemental Table S2).

**Figure 3.**
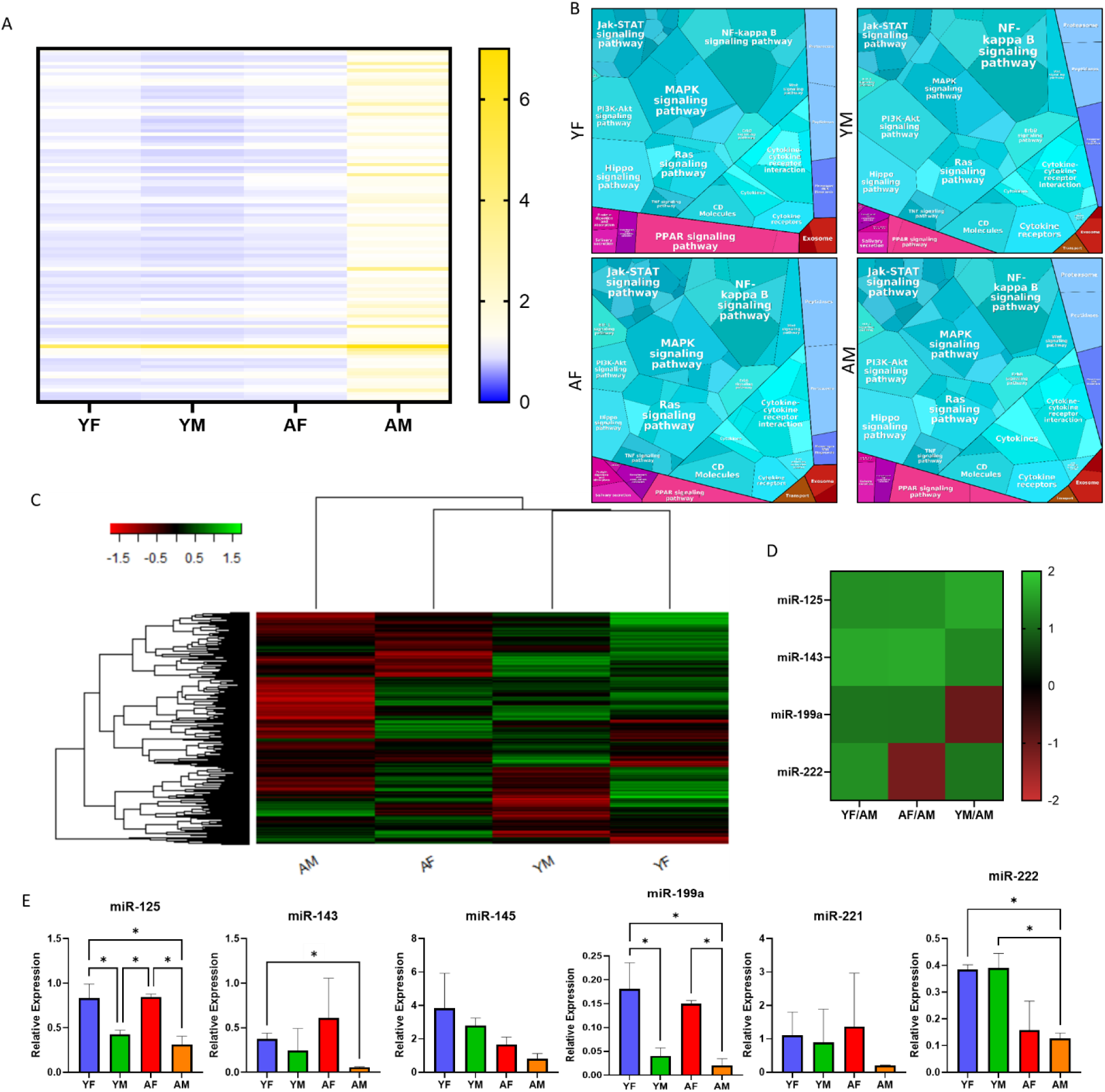
Left Ventricular Vesicles (LVVs) Affect a Wide Variety of Aging and MI-Related Pathways Depending on Age and Sex. (A) Heat maps showing the cytokines detected in each cohort relative to the internal positive control. For read data, see Supplemental Table S1. (B) Proteomaps showing the KEGG pathways affected in response to the cytokines in each cohort. (C) Heatmap showing the full miRNA profiling for each biological replicate of each cohort. (D) Nanostring results for the 6 identified exosomal miRNAs from literature, for each cohort relative to AM. (E) PCR results for the identified miRNAs. Data are presented as the mean ± standard deviation. * p < 0.05, assessed by one-way ANOVA with Tukey’s post-hoc for (E).

### Identification of miRNA Content

miRNA profiling via Nanostring analysis revealed highly upregulated exosomal miRNA populations in both young groups relative to both aged groups (Figure 3C, Supplemental Figure S9). Interestingly, many of the miRNAs upregulated in YF, YM, and AF LVVs were downregulated in the AM group. From the over 800 miRNAs profiled, six were identified as both exosomal and cardioprotective from literature^31–34^, although the activities of these miRNAs have been primarily characterized for cardiomyocytes. Of these six, five were upregulated in other groups relative to AMs (Figure 3D). RT-qPCR revealed that four were significantly increased in at least one comparison against AM (Figure 3E), with trends being consistent with the Nanostring data. Interestingly, miR-125 and miR-199a were elevated in both female groups compared to both male groups, while miR-143 was decreased only in AM LVVs. Additionally, miR-222 was elevated in young groups compared to aged. Both miR-145 and miR-221 showed no significant differences between any groups, which was unexpected as these miRNAs are often considered as conjugated units with miR-143 and miR-222, respectively.

### Features of LVVs are Recapitulated in Mouse Models

Preliminary results had shown that LVV effects were mostly consistent between human and rat models (Figure 2F, Supplemental Figure S5), so a more controlled mouse study was conducted to further validate these results and account for the biological variability and difficulty of obtaining additional human samples. Mice (n=6) were similarly categorized as YF, YM, AF, or AM (Supplemental Table S3). The collagen contraction assay using mouse LVVs (mLVVs) and mouse CFs (mCFs) showed a similar trend to that observed from human samples, but both young groups showed no improvement compared to the untreated control, and the AM group showed no significant increase in contraction in the same comparison (Figure 4A). However, the AF-treated group showed a significant decrease in contraction compared to the control, and YF, YM, and AF groups showed a significant decrease in contraction compared to the AM group. The results from the wound healing assay followed this trend, and YM and AF, but not YF or AM, were significantly different from the control (Figure 2B). However, once again the YF, YM, and AF groups showed significantly increased wound healing compared to the AM group. These results echo the preliminary results obtained from rat models (Supplemental Figure S5).

**Figure 4.**
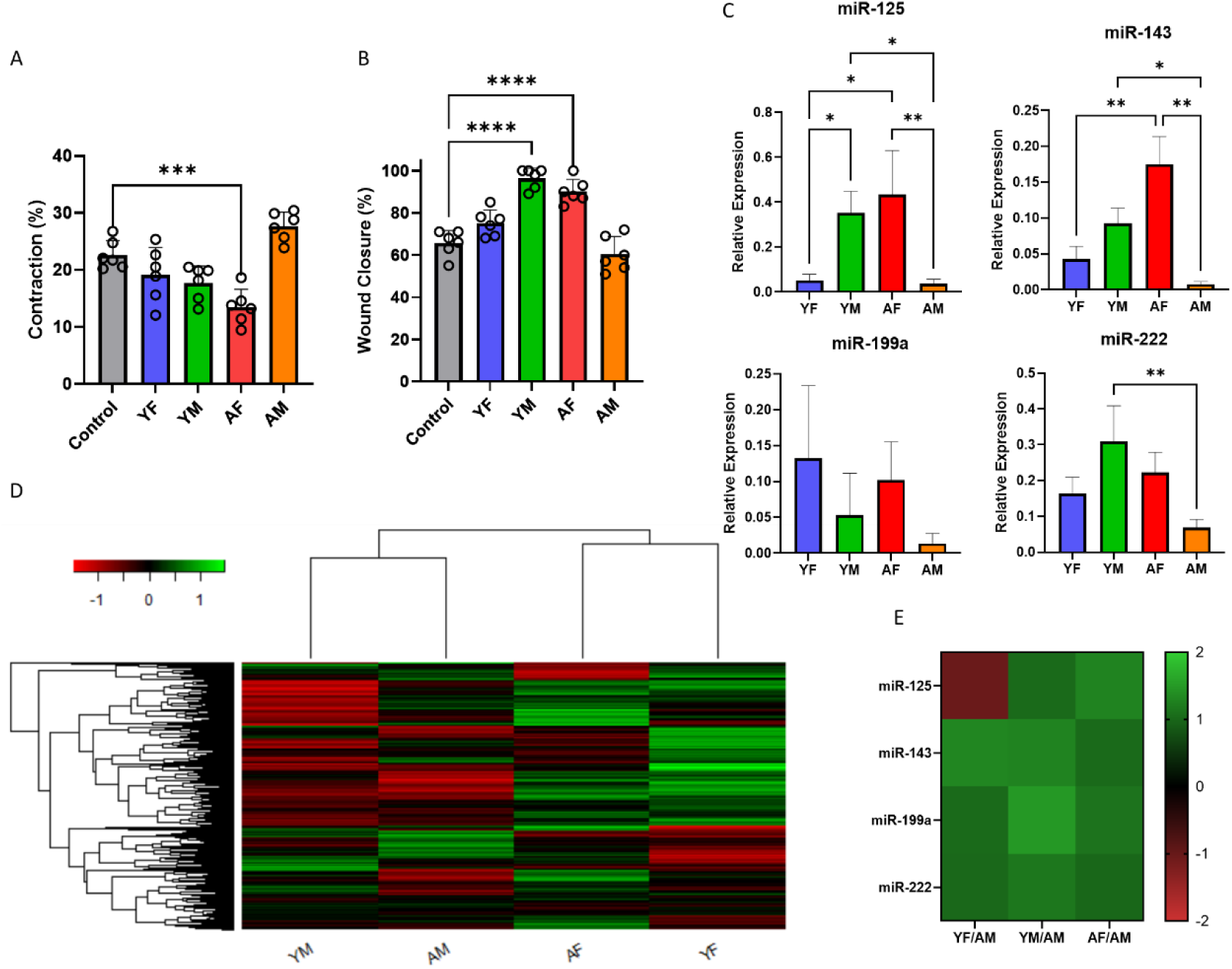
Validation of Trends in Human LVVs in Mouse Cardiac Model EVs. (A) Gel contraction over 48 h for collagen hydrogels, as a percentage of initial gel diameter (n = 6 biological replicates for each cohort, for 2 independent repetitions). (B) Wound closure over 60 h, as a percentage of initial wound area (n = 6 biological replicates for each cohort, for 2 independent repetitions). (C) PCR results for the 4 miRNAs selected from human LVVs (n = 3 biological replicates, each pooled from 2 separate hearts’ miRNA, for 2 independent repetitions). (D) Heatmap showing the full miRNA profiling for each replicate used for PCR (each replicate is from 2 separate hearts’ isolated miRNA) and (E) a heatmap for each cohort for the 4 selected miRNA targets (from the data presented in the full profiling). Data are presented as the mean ± standard deviation. Bar graphs represent biological replicates, with individual replicates overlaid as a dot plot. * p < 0.05, ** p < 0.01, *** p < 0.001, **** p < 0.0005, assessed by one-way ANOVA with Tukey’s post-hoc for (A), (B), and (C).

Also assessed in mLVVs was the relative expression of the target miRNAs, excluding those which showed no significance in human samples. First these four targets were measured by RT-qPCR (Figure 4C). Three of the four targets showed significantly increased expression in at least one group compared to the AM cohort, with miR-199a demonstrating no significant difference between any groups. Additionally, the trends in miRNA expression slightly differed between mouse and human LVVs. This difference can likely be attributed to innate differences in mouse and human physiology and native response to cardiac insult, which may be fundamentally different^35^. Nevertheless, the AM group still showed the lowest expression of all four target miRNAs, and that, with the decreased in some miRNA expression observed from the YF group, is co-concurrent with mitigated benefits in contraction and wound healing. Full miRNA profiling of the mLVVs was also performed (Figure 4D). There was a higher overall upregulation of miRNAs in both female groups compared to both male groups. Interestingly, among upregulated miRNAs there was notable overlap between the YM and AF groups, although this overlap also commonly includes the YF or AM groups. The 4 selected target miRNAs pulled from the total profiling were mostly consistent with the RT-qPCR results, although for the YF group miR-125 showed downregulation compared to the AM group (Figure 4E).

### miRNAs Partially Recapitulate LVV Effects

To study the effect of miRNAs on wound closure, cells were treated with miR-125, miR-143, miR-199a, or miR-222 mimics, or a cocktail containing all four. No significant differences were observed from treatment with individual miRNAs, although some increase in wound healing was detected after treatment with miR-125, while the miRNA cocktail-treated groups demonstrated up to three-fold greater wound closure compared to the scramble siRNA-treated control over the culture period (Figure 5A). At 48 h, the miR cocktail-treated cells had healed significantly more than scramble siRNA treated control cells. Staining of the cells (Figure 5B) revealed a significant decrease in α-SMA expression of the treated cells compared to control (Figure 5C) with no change in vimentin expression (Figure 5D), as was observed with LVV treatment. Quantification of Live/Dead staining (Figure 5E) and BrdU staining (Figure 5F) of hCFs subjected to MI-like conditions (3 h hypoxia) showed about a 2.5-fold increase in the ratio of live cells to dead cells after treatment with miRNA cocktail compared to control cells (Figure 5G), while relative BrdU expression was similar between both groups (Figure 5H).

**Figure 5.**
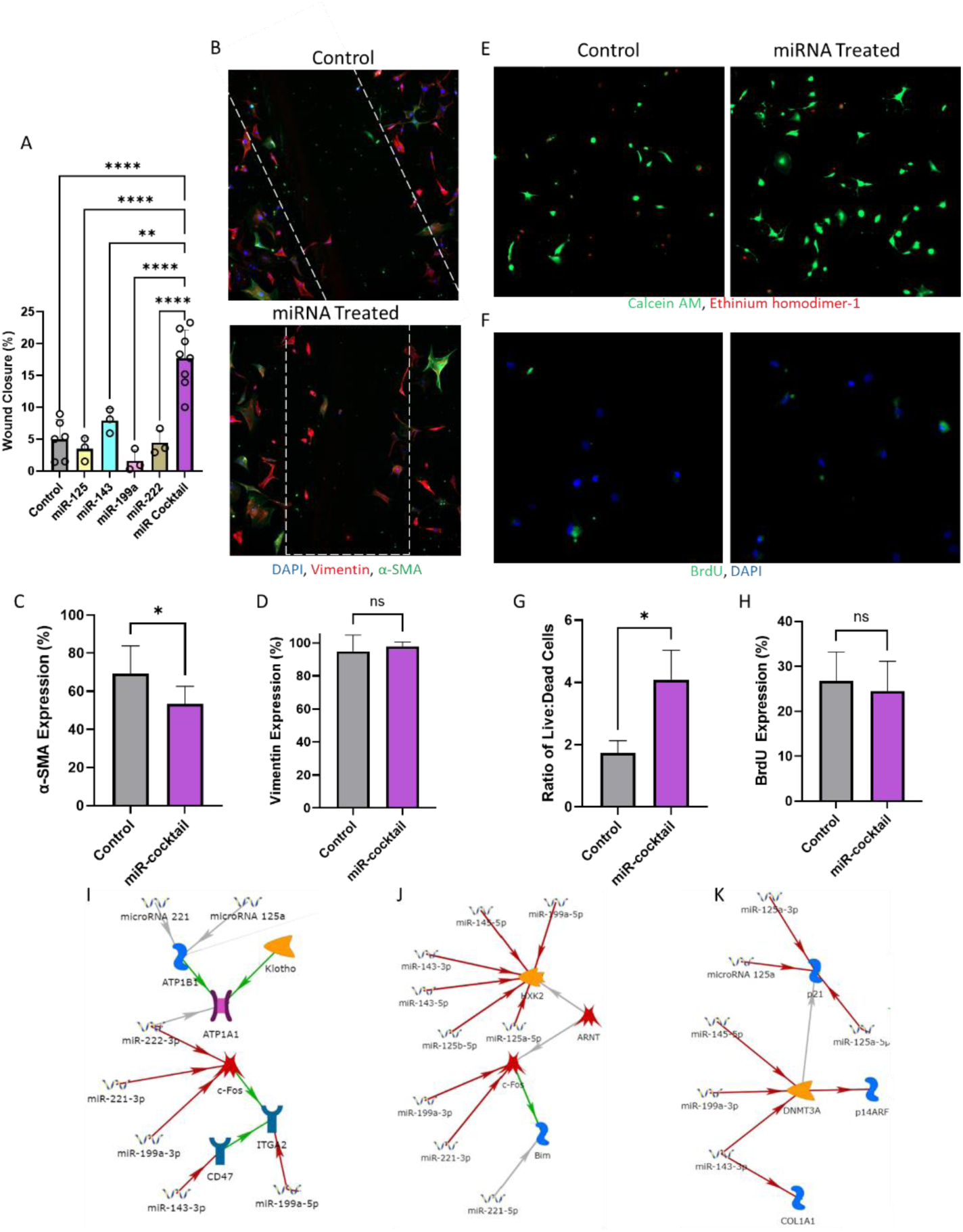
miRNA Treatment Partially Recapitulates LVV Effect *in vitro*. (A) Final timepoint wound closure percentage of control and miRNA-treated groups (n ≥ 3, 2 technical replicates per sample, 3 images per replicate), with individual replicates shown as a dot plot overlay. (B) Immunostaining showing expression of DAPI (blue), α-SMA (purple), and vimentin (green) for control or miRNA-treated hCFs post-healing. Quantification of local expression of α-SMA (C) and vimentin (D) as a percentage of total cells observed. Representative images showing (E) live/dead stain or (F) BrdU stain following MI-like hypoxia treatment of control or miRNA-treated cells (n ≥ 3, 3 images per sample). Quantification of (G) live/dead or (H) BrdU assay. (I-J) Results of MetaCore pathways analysis for the identified target miRNA. Data are presented as the mean ± standard deviation. * p < 0.05, assessed by ANOVA with Tukey’s post-hoc for (A), and Student’s t-test with Welch’s correction for (C), (D), (G), and (H).

### Target miRNAs Regulate Fibrosis-Related Pathways

To assess potential convergence on fibrosis-related pathways of interest, MetaCore pathway analysis software was utilized to build networks for the selected target miRNAs. These networks identified several pathways of interest that were regulated by two or more of the selected targets. Of interest was the regulation of c-Fos by miR-199a-3p, miR-221, and, interestingly, miR-222, and downstream regulation of integrin subunit alpha 2 (ITGA2) by miR-199a-5p, which was also regulated by miR-143-3p through CD47 (Figure 5I). This same pathway also showed regulation downstream of Klotho by miR-221 and miR-125a through ATPase Na+/K+ transporting subunit beta 1 (ATP1B1), and directly by miR-222. Also of interest was regulation of hexokinase 2 (HXK2) by miR-125a-5p, miR-125b-5p, miR-143-3p, miR-143-5p, miR-145-5p, and miR-199a-5p, again in parallel to c-Fos regulation (Figure 5J). Finally, DNA methyltransferase 3 alpha (DNMT3A) regulation by miR-143-3p, miR-145-5p, and miR-199a-3p, as well as downstream regulation of p21 by both forward and reverse strands of miR-125a (Figure 5K). Also interesting was the relationship between miR-143-3p and collagen type 1 alpha chain 1 (COL1A1) expression. Full maps with all nodes were also recorded for each identified pathway (Supplemental Figure S10).

## DISCUSSION

In this study, we isolated and characterized the matrix-bound vesicles in left ventricular tissues of young (19-40 years old) and aged (50-63 years old) male and female human donors and assessed their effect on cardiac fibroblast transdifferentiation through measurement of contractility, wound healing, and α-SMA expression. Interestingly, we found differences in size distribution, uptake, and cytokine and miRNA profiles between aged and young LVVs, as well as male and female LVVs. LVVs from aged hearts were smaller in size than those from young and were taken up more rapidly by cells. Additionally, LVVs from all cohorts expressed common exosome markers, showing that at least some of the isolated LVVs were exosomes. Cytokine content was higher in LVVs from aged tissues compared to young, and in males compared to females, with the highest content observed in AM LVVs. Conversely, the lowest miRNA content was observed in AM LVVs. While LVVs from females contained more miR-125 and miR-199a than LVVs from males, those from young tissues contained more miR-222 than aged. Remarkably, hCFs embedded in collagen gels showed increased contraction upon treatment with AM LVVs compared to untreated controls and decreased contraction when treated with LVVs from other groups. Similarly, *in vitro* scratch assay showed decreased wound closure in AM LVV treated cells compared to untreated controls and increased closure in cells treated with other LVVs groups in both human and murine cell lines. Immunostaining of cells post-scratch assay showed higher α-SMA expression in AM LVV-treated cells than the control, and lower expression in other groups. Repetition of the scratch assay using miRNA treatment instead of LVVs showed that miRNA cocktail treatment, but not individual miRNAs, could recapitulate these effects. Remarkably, treatment with this cocktail showed a protective effect on hCFs subjected to MI-like conditions, significantly decreasing cell death without significantly affecting proliferation. These miRNAs were then found to be involved in the regulation of some components of fibrosis-related pathways at different points, suggesting that these miRNAs and others may work together on parallel pathways in order to regulate fibrotic signaling. These results show for the first time that matrix bound EVs in the left ventricle change in size, content, and bioactivity in an age-and sex-dependent manner, and that AM LVVs may play a novel and pivotal role in chronic pro-fibrotic cardiac signaling.

In this study we showed for the first time the presence of exosome-like EVs in human left ventricles and characterized their size and characteristics as a function of age and sex. Interestingly, although EVs from all cohorts were in the size range of exosomes (30-200 nm)^36^, both young groups showed greater size and dispersity than both aged groups despite similar concentrations and measured quantity isolated. These data suggest that there is a physical difference between the LVVs present in aged ECM compared to young ECM. It is established that cell uptake of vesicles is size-dependent for exosomes^37^, so this may suggest a need for more rapid uptake of these exosomes for expedited response in older, more “at risk” hearts. Alternatively, the production of smaller vesicles may result from increased vesicle specialization in aged tissue^23, 27^, although this does not explain the increased cytokine content observed in the AM LVVs. In either case, these data suggest substantial differences in exosome uptake and secretion mechanics between young and aged hearts. While the smaller, aged tissue EVs were taken up more rapidly and to a greater degree than young tissue EVs, anti-fibrotic effects were mostly conferred by young LVVs. This suggests that the miRNA-mediated effects conferred by young EVs are not intended for a rapid response or may not be highly dose-dependent. Alternatively, the detrimental effects observed from AM LVVs might result from their high cytokine content. These findings suggest a different core paracrine response to cardiac injury between young and aged, and male and female myocardium which has not been previously described in literature.

A general upregulation of most assessed cytokines was observed in AM LVVs compared to other cohorts, as was expected based on available data for sex^38^ and age^39^ dependence of cytokine profiles. Cytokines related to pro-fibrotic processes post-MI and other detrimental cardiac processes were present in greater amounts in AM LVVs compared to other cohorts. While the AM cohort demonstrated the highest upregulation of inflammatory cytokines, both aged groups showed some upregulation of cytokines such as VCAM-1 and IL-1β (Supplemental Table S1), which are involved in further injury or fibrosis post-MI^40, 41^. These data suggest that AM LVVs may participate in microenvironment-driven inflammaging, which has been suggested as a major contributor to cardiac fibrosis^42^. Some other cytokines elevated in AM LVVs, such as IFN-γ, MMP9, and myeloperoxidase, which are involved in cardiometabolic dysfunction^43^ and fibrotic remodeling post-MI^44–46^, also suggest that this contribution is greater from AM LVVs, and that AM LVVs may more directly contribute to long-term cardiac damage from the microenvironment. Interestingly, AM LVVs also demonstrated greater levels of cardioprotective cytokines than other cohorts. This may be due to endogenous ischemic preconditioning, which has been suggested to be mediated by atypical cytokine interactions^47–50^. Since neither AM subject suffered from CVD or other diseases which may affect the heart microenvironment or died from a heart-related cause, the observed high levels of cytokines may be a result of ongoing endogenous preconditioning. In fact, the AM group demonstrated increased GM-CSF and GDF-15, SDF-1α, and TNF-α, which have been identified as major signaling molecules in ischemic preconditioning^48, 51^. Overall, AM demonstrated the greatest expression of both damaging and protective cytokines.

A different trend was observed from miRNA analysis of six cardioprotective miRNA, with AM consistently demonstrating the lowest expression of all miRNAs assayed. This was apparent in the full miRNA profiling, where YF LVVs show the most consistent upregulation of miRNAs and AM LVVs show the most consistent downregulation of miRNAs, for all 800+ miRNAs assessed. Based on this trend, potential miRNA targets were identified by comparison of expression levels relative to AM expression to find the largest upregulation. First, targets were selected from literature and validated using the profiling data, then quantified using RT-qPCR. The targets were as follows: miR-125, shown to protect against ischemic and reperfusion injury^32, 52^, was significantly increased in female tissue LVVs in both aged and young subjects. miR-199a, a cell survival promoter and key regulator of the endothelial nitric oxide pathway^31, 32^, was significantly increased in AF LVVs compared to both male groups. These data suggest a shift in miRNA production may be integral to differences in response to cardiac event between males and females. While miR-143, implicated in regulation of cardiac regeneration and protective against carotid injury^53, 54^, was significantly increased in YF LVVs compared to AM, this may result from a combination of age and sex differences. Additionally, miR-222 was age-dependent, while miR-145 and miR-221 showed no significant difference between any groups. This is an interesting result particularly because miR-143 and miR-145, and miR-221 and miR-222 are often considered as conjugated pairs rather than individual miRNAs^32^.

Cell assays were selected to determine relative transdifferentiation of CF samples treated with the same concentration of LVVs from each group. These provided metrics of contractility, which is enhanced in MFs^40, 55^, proliferation and proliferative wound healing, which are decreased in MFs^55, 56^, and expression of characteristic MF marker α-smooth muscle actin (α-SMA)^40^. These data show that LVVs from the AM cohort tended to promote MF-like behavior from cells in both 2D and 3D culture. This is an interesting result, as existing literature suggests that the application of ECM in general tends to promote a reparative, anti-fibrotic microenvironment in the myocardium^17–19^. While these anti-fibrotic effects are still observed from AF LVVs and both young LVV groups, this is not the case for AM LVVs, suggesting that AM LVVs contain distinctly pro-fibrotic factors. This observation aligns with expected effects from heart tissue subjected to inflammaging effects^42^. Furthermore, this is supported by additional preliminary data on rat cardiac fibroblasts as well as well-controlled trials with mouse cardiac fibroblasts. In both cases, both the YM and AF groups demonstrated beneficial effects consistent with the human trials, while the AM group demonstrated pro-fibrotic effects to a comparable degree as direct TGF-β treatment in the rat model. In the mouse model, the AM group was significantly detrimental compared to all other treatment groups, further suggesting that some aspect of AM LVVs is inducing pro-fibrotic effects in direct opposition to other cohorts in this study. To further compare LVVs with human biofluid EVs, we performed WHA on EVs isolated from plasma from YF, YM, AF, and AM individuals (Supplemental Figure S11). While this assay was performed with iPSC-derived CFs, the results were consistent with the hCF WHA results and showed that LVVs demonstrated significantly increased efficacy in wound healing compared to the plasma EVs. Additional NS analysis was performed on the plasma EVs, and comparison to the LVV NS results showed that the miRNA targets identified were mostly localized to the tissue-bound LVVs, although some of the targets were found to be elevated in the female group plasma EVs. These results further demonstrate the validity of identifying miRNA targets from the tissue-bound LVVs instead of plasma EVs for the development of therapeutic strategies.

Further investigation of AM LVV factors may implicate novel targets for therapeutic intervention of aging-related inflammatory pathways which stimulate cardiac fibrosis. Remarkably, however, the AF, YM, and YF LVVs exhibited anti-fibrotic effects, similar to what has been observed from treatment with decellularized ECM^17–19^. Of these, the YF group demonstrated the greatest reduction in contractility and α-SMA expression, while AF LVVs promoted the greatest increase in wound healing. This is an interesting result, as LVVs from the female subjects displayed the greatest anti-fibrotic behavior overall, consistent with clinical outcomes for the onset of fibrosis. Together, these findings suggest that the beneficial effects of LVVs are sex and age-dependent, and that LVVs can recapitulate the beneficial effects of ECM or introduce pro-fibrotic factors depending on these conditions. Further investigation of these differences will help elucidate the mechanistic reason for the clinically observed importance of sex and age in the onset of cardiac fibrosis.

To better understand these mechanisms, we attempted to recapitulate the effects of LVV treatment by transfecting hCFs with the identified miRNAs of interest: miR-125, miR-143, miR-199a, and miR-222. While transfection with individual miRNAs did not yield significant results, treatment with a combination of all four miRNAs enhanced wound healing and survivability and decreased transdifferentiation of hCFs, similarly to treatment with LVVs. Most interestingly, this miRNA cocktail exhibited these same effects under MI-mimicking conditions. Treatment more than doubled cell survivability after 3 hours of MI compared to the control while cell proliferation was nearly unchanged, suggesting that the synergistic effects of these miRNAs not only reduce damage from CVD events but also promote survival and “reparative” signaling. This is an exciting result, and indicates that novel synergistic effects of exosomal miRNAs can both inhibit the onset of cardiac fibrosis and protect cells from MI-induced cell death through endogenous pathways, although further targets must be identified to better recapitulate the full effects of LVV treatment. Similar to how the application of mesenchymal stem cell exosomes can recapitulate the cardioprotective effects of the source cells^14, 57^, the isolation of key cardioprotective agents, such as exosomal miRNA combinations, from the ECM may allow for enhanced treatment options. By investigating key differences observed between the AM LVVs and other cohorts and common factors between AF, YM, and YF LVVs, the mysteries of the myocardial microenvironment in MI and cardiac fibrosis will be elucidated.

In that vein, given the success of treatment with only four miRNAs, we have identified 37 additional targets for either application or inhibition based off the miRNA profiling data (Supplemental Table S4). Specifically, 14 targets are exosomal miRNAs that display similar elevated expression against the AM group to the selected targets, while the remaining 23 are miRNAs that are elevated in the AM group compared to others. While above we have established the benefits of utilizing a synergistic cocktail of miRNAs, other single-miRNA studies have suggested that some miRNAs may also be driving factors behind the pro-inflammatory signaling and other microenvironment changes observed in the heart^58^ and profibrotic changes observed in other organs^59^.

Beyond our own dataset, we also suggest that this data can be used to identify downstream druggable targets for both study and therapeutic intervention that are not directly implicated in this dataset. This is due to the converging of target miRNAs on several pathways which have been implicated in fibrotic signaling, such as CD47^60^, ITGA2^61, 62^, c-Fos^63, 64^, HXK2^65, 66^, Klotho^67^, p21^68, 69^, and DNMT3a^70^. One particularly interesting connection is that the TGF-β-mediated effects of CD47 and ITGA2 appear downstream of c-Fos, which is known to be involved in AngII-mediated fibrosis^63^ and can be regulated by some miRNAs to impact the onset of TGF-β-mediated fibrosis effects^64^ in conjunction with AngII-mediated fibrotic signaling around p21^68^. Additionally, the epigenetic changes implicated in fibrosis for both p21 and DNMT3A related pathways appear to be mediated by miRNAs or some other paracrine pathway^69–71^. Further overlap occurs in the regulation of HXK2 and Klotho, which have both been shown to mediate fibrosis at least in part through regulation of the TGF-β-Wnt axis^65, 67^, a highly sought after regulatory pathway which is notoriously difficult to regulate. These points all demonstrate that the selected miRNAs, though limited, already demonstrate multi-point regulation of key fibrosis-mediating pathways and suggest reinforce the synergistic effects of this endogenous miRNA cocktail. By expanding our knowledgebase of which miRNAs may be contributors to this synergistic, anti-fibrotic signaling, it is possible to further refine which pathways these miRNAs may be acting on in order to better map out the process by which chronic cardiac fibrosis is mitigated endogenously to better develop targeted therapeutic strategies.

In this study we showed, for the first time, the presence of matrix bound exosome-like EVs in the human left ventricle, characterized the physical properties and cytokine and miRNA contents of these LVVs as a function of age and sex, and investigated the effects of these LVVs on cardiac fibroblast transdifferentiation. While recent studies have identified the therapeutic effects of young plasma-derived exosomes^72, 73^, this study is among the first to directly compare exosomes from young and aged subjects, an identified gap in knowledge regarding exosome studies^4, 30^, and is the first to do so with exosomes derived from cardiac tissue. Additionally, this study contributes to understanding of exosome behavior in healthy hearts, which is understudied compared to the knowledgebase for exosomes from hospitalized subjects^30^. Furthermore, this study reveals previously undescribed synergistic effects of exosomal miRNAs in the progression of cardiac fibrosis and MI-induced damage to cardiac fibroblasts. This study expands the knowledgebase of changes in exosome behavior related to cardiac health during aging and contributes to the identification of factors involved in cardiac fibrosis and MI. These factors can be subsequently developed as a cell-free class of endogenous therapeutics, or otherwise identify highly specific therapeutic targets, that can mitigate or prevent these phenomena.

In conclusion, functional exosomes can be found embedded within the ECM of human left ventricular tissue, differing from exosomes traditionally isolated from biofluids. These novel left ventricular vesicles, or LVVs, contain varying quantities of cytokines and miRNA depending on sex and age, with LVVs from aged male subjects showing signs of inflammaging, ischemic preconditioning, and decreased cardioprotective miRNAs. Treatment of fibroblasts revealed that LVVs from aged males tend to promote a pro-fibrotic response, whereas LVVs from females and young males promote an anti-fibrotic response. These data suggest that ECM-embedded vesicles play a crucial role in response to cardiac injury and may be responsible for some cardioprotective effects observed with ECM treatment. Furthermore, these effects were also observed upon treatment of hCFs with a cocktail of cardioprotective miRNAs identified from the LVVs: miR-125, miR-143, miR-199a, and miR-222. This cocktail promoted cell survival under MI-like conditions *in vivo* while partially recapitulating the wound healing effects observed from LVVs. This suggests that miRNAs can mimic the effect of exosomes and possibly be used as therapeutic agents for cardiac fibrosis. However, it should be noted that physical and content characterization of the exosome populations was limited by the small number of human biological replicates available. With only up to three human subjects from each cohort, it is possible that significant effects were overlooked in this study and internal variability was exacerbated. Nevertheless, further study of these interactions will enhance understanding of the mechanisms by which cardiac injury response occurs and provide novel means for intervention.

## METHODS & MATERIALS

### Tissue Preparation

Human heart tissue was collected from donors whose hearts were deemed unsuitable for transplantation through the Indiana Donor Network. IRB approval was waived, as no identifying information was provided by the Indiana Donor Network. All tissue collection was performed in accordance with the declaration of Helsinki. Subjects were selected such that cardiac event or cardiovascular disease was not the primary cause of death. Tissue samples consisted of young female (YF, N = 2), young male (YM, N =3), aged female (AF, N = 3), and aged male (AM, N = 3) subjects, where subjects at or over 50 years old were considered “aged”, and those at or below 40 years old were considered “young”, with all but one YM sample falling below 30 years old (Supplemental Table S5). Samples were stored at −80 °C prior to sectioning. While still frozen, extraneous fat and connective tissue were excised. Tissue was thawed in sterile PBS at 4 °C and sectioned. Sections with approximately the same surface area and thickness (<300µm) were processed.

Mouse heart tissue was collected from C57BL/6J mice (The Jackson Laboratory) according to IACUC guidelines (protocol number: 18-05-4687) with the approval of the University of Notre Dame. Male and female mice were categorized as young (16 weeks old) or aged (72 weeks old), corresponding to the ages of the collected human samples (Supplementary Table S1), with each group having n = 12 samples per group (N = 48 total mice). Mice were euthanized via CO_2_ and the whole mouse heart and other tissues were immediately harvested. All mice showed no cardiovascular abnormalities upon death or tissue isolation. Following collection, the left ventricle of the hearts were isolated and immediately processed.

### Decellularization

Decellularization and digestion were performed in accordance with current standards for maintaining EV integrity^74, 75^. Tissue sections were agitated in a solution containing peracetic acid (Sigma Aldrich, USA) (0.1%) and ethanol (Sigma Aldrich) (4%)) at 200 rpm for 2 h, then in phosphate buffered saline (PBS) at 200 rpm for 2 h, and then again in peracetic acid/ethanol solution at 200 rpm for 16 h. Decellularized matrix sections were then washed extensively in PBS and sterile water, blotted on a tissue paper, and frozen at −80 °C.

### Decellularized Matrix Digestion

Frozen heart matrices were lyophilized overnight and ground into a powder using liquid nitrogen and pre-chilled mortar and pestle. ECM powder (200 mg) was then transferred to 1.5 mL microcentrifuge tubes, suspended in 1 mL of digestion buffer containing 0.1 mg/mL collagenase type II (Corning), 50 mM Tris buffer (Sigma Aldrich), 5 mM CaCl_2_ (Amresco), and 200 mM of NaCl (Sigma Aldrich), and was mixed vigorously to ensure complete resuspension. The mixture was stored statically at room temperature (RT) for 24 h or until few or no solid particles could be observed in the solution, with brief remixing every 6-8 h.

### Vesicle Extraction and Isolation

Digested matrix solution was centrifuged three times at 500g for 10 min, 2500g for 20 min, and 10,000g for 30 min, and the pellet discarded after each centrifugation step to remove any remaining insoluble matrix remnants. The final supernatant was centrifuged at 100,000g at 4°C for 70 min using an ultracentrifuge (Optima MAX-XP Tabletop Ultracentrifuge, Beckman Coulter). The pellet was either used immediately or stored dry at −80°C.

### Transmission Electron Microscopy

Single pellets were fixed in 2.5% glutaraldehyde at RT in the dark, then loaded onto plasma-cleaned Formvar/carbon-coated copper 200 mesh grids (Polysciences) and negative-stained with Vanadium staining solution (Abcam, ab172780). Samples were imaged at 80 kV with a TEM (JEOL 2011, Japan).

### Nanoparticle Tracking Analysis

Single pellets were resuspended in 1mL of sterile, particle-free PBS and measured using a NanoSight NS300 machine (Malvern Panalytical) and NTA software version 3.2.16. This method obtains the hemodynamic diameter and concentration of nanoparticles with diameters from 10-1000 nm in solution via Brownian motion analysis. Samples were kept at 4 °C until measurement, and measurements were taken at RT.

### Western Blot

To retain maximum protein blot clarity, decellularization and matrix digestion were performed at 4°C. The pellets were lysed in RIPA buffer containing 1% proteinase inhibitor cocktail (Brand, Country) at 4°C for 30 minutes, then protein concentration was assessed via bicinchoninic acid (BCA) assay (Pierce Chemical). Equal amounts of protein were separated by 12% SDS-PAGE and transferred to blotting membranes, which were incubated overnight at 4°C with the rabbit polyclonal primary antibodies against CD9 (Abcam, ab223052), CD63 (Abcam, ab216130), TSG101 (Abcam, ab30871), Syntenin-1 (Abcam, ab19903), Vimentin (Abcam, ab137321), Connexin 43 (CX43, Abcam, ab11370), and VE-Cadherin (Abcam, ab33168) at (1:1000) dilutions, and against GRP94 (Abcam, ab3674) at 1:2000 dilution, then for 1 h at RT with HRP-conjugated goat anti-rabbit secondary antibody (Abcam, ab205718). Membranes were then exposed to a chemiluminescent substrate (Clarity ECL, Bio-Rad) and imaged using a ChemiDoc-It2 imager (UVP, Analytik Jena) equipped with VisionWorks software. Images were processed using ImageJ (NIH).

### Cell Culture

Human cardiac fibroblasts (hCFs) were obtained from Cell Applications (USA) at passage 1, and cultured in Dulbecco’s Modified Eagle Medium (DMEM) (Thermo Fisher) supplemented with 10% fetal bovine serum (FBS) (Gibco), 1% penicillin/streptomycin (P/S) (Life Technologies), henceforth called DMEM Complete, and 3 µM SD208, a TGF-β receptor I kinase inhibitor (Sigma Aldrich). Cells were cultured with SD208 supplement to inhibit transdifferentiation, and then used between passage 4 and 10 without SD208. Mouse cardiac fibroblasts (mCFs) were obtained from iX Cells Biotech (USA) at passage 0, and were cultured under the same conditions as hCFs.

### Cell Uptake of EVs

hCFs were seeded in a 24-well plate at 50,000 cells/well to allow for imaging of small cell clusters without compromising cell viability. Three wells were seeded for each biological replicate of LVVs and an empty control. LVVs were stained with ExoGlow (System Biosciences), according to the manufacturer’s protocol. Briefly, LVV content was quantified with the bicinchoninic acid (BCA) gold protein quantification assay (Thermo Fisher Scientific), and 25 µg of LVVs was obtained from each sample and resuspended in 12 µL of provided reaction buffer. After, 2 µL of stain was added and allowed to react for 30 min at RT. Stained LVVs or an empty control were then isolated in a provided gradient column and resuspended in 2mL of DMEM with 1% P/S, according to the manufacturer’s protocol, for a final concentration of 12.5 µg/mL. During this, hCFs were incubated with Cell Tracker Green (Thermo Fisher) in PBS for 30 min at 37°C. Stained cells were incubated in DMEM with 1% P/S with one group of LVVs or the control at 37°C for 24 h, and imaged at 3, 8, 16, and 24 h of incubation. Before imaging, the conditioned media from each well was moved to a sterile container and the cells were washed with PBS. Cells were imaged in PBS, and the removed media was replaced after imaging.

### Gel Contraction Assay

Collagen solution (1.5 mg/mL) was prepared by mixing rat tail collagen (9.33 mg/mL, Corning), 10x PBS, deionized (DI) water, and DMEM with 1% P/S at 2:1:8:3 ratio in a final volume of 250 µL. Immediately before seeding, pH was adjusted to 7.4 with 1 M NaOH. All steps up to the addition of cells were performed on ice to prevent premature gelation. Cells were washed with PBS, detached from flasks using trypsin-EDTA (0.25%), and then resuspended at 1.5 million cells/mL in DMEM with 1% P/S. Cell suspension was mixed thoroughly with the collagen solution at 1:1 ratio. The mixture was transferred into a 24-well plate (300 µL/well) and incubated at 37 °C for 2 h to allow for gel formation. In addition to five cell-encapsulated gels, one cell-free gel (loaded with FBS-free DMEM Complete without cells, henceforth gel control) was included. Gels were then incubated in FBS-free DMEM Complete supplemented with 12.5 µg/mL LVVs from one group, or a PBS blank containing no LVVs (control). The gel control was fed with control media. Images were taken every 12 h for 48 h and the diameter of the gel was measured along two sets of orthogonal axes. This experiment was repeated twice (3 repetitions total) for each of the biological replicates for all cohorts.

### Wound Healing Assay

Cells were seeded onto a 24-well plate at a density of 2 x 10^5^ cells per well and allowed to grow to >90% confluency. Once confluent, cells were washed with PBS and subjected to a vertical wound by gently dragging a 1000 µL pipette tip across the monolayer. Wells were assessed under the microscope to ensure successful and consistent wounding. Typical wound width was approximately 600 µm. The cells were then incubated in FBS-free DMEM Complete supplemented with 12.5 µg/mL LVVs or a PBS blank containing no LVVs (control). The wounds were then imaged immediately and every subsequent 24 h for 96 h. Cells were incubated at 37 °C between imaging, and media was replaced after 48 h. Wound healing was assessed using ImageJ by percent reduction in wound width in three locations over time. This experiment was performed twice (2 repetitions total) for each biological replicate for all cohorts.

### Immunostaining

At 96 h post-wounding, cells were washed with PBS and incubated in 4% paraformaldehyde for 15 min, then in 0.1% Triton X-100 for 30 min, and then in 10% goat serum for 2 h, all at RT and with PBS washes after each step. Cells were next incubated with rabbit anti-vimentin (Abcam) and mouse anti-α-SMA (Abcam) primary antibodies (dilution: 1:100 in 5% goat serum) at 4 °C overnight. The cells were then washed and incubated with Alexa Fluor 647-labelled anti-rabbit IgG and Alexa Fluor 488-labelled anti-mouse IgG secondary antibodies (dilution: 1:200 in 5% goat serum) at 4°C for 6 h. Finally, the cells were incubated with DAPI (dilution: 1:1000 in PBS) for 15 minutes at RT and imaged with a fluorescent microscope (Axio Observer.Z1, Zeiss).

### Profiling of Cytokines

To remove any residual extraneous proteins, samples were purified using the CD9 Exo-Flow Capture Kit (System Biosciences) using the manufacturer-provided protocol. Briefly, the pellet was resuspended in a solution of biotin-conjugated CD9 antibody and streptavidin-coupled magnetic beads overnight at 4 °C. LVVs were isolated magnetically and washed, then eluted from the magnetic beads. The LVV solution was worked up to 1% Triton X-100 and left at 4 °C overnight. The lysed LVV solution was assessed using the Proteome Profiler Human XL Cytokine Array Kit (R&D Systems) as described previously^76, 77^, for detection of 111 cytokines (Supplemental Figure S8, Supplemental Table S6). Briefly, nitrocellulose membranes with the immobilized antibodies against 111 cytokines were blocked according to manufacturer’s instructions and incubated overnight at 4 °C with equal concentrations of proteins, determined by BCA assay, from each sample. Membranes were then washed and incubated with antibody cocktail solution for 1 h, with streptavidin-horseradish peroxidase (HRP) for 30 min, and with the Chemiluminescence reagent mix for 1 min. Membranes were then imaged with a biomolecular imager (ImageQuant LAS4000, GE Healthcare) using X-ray exposure for 5-10 min. Relative cytokine content was determined by blot intensity analysis in ImageJ.

Gene Ontology Analysis (GOA) was performed on the obtained relative expression data. Comprehensive analysis was performed using an online database via PANTHER Gene Ontology classification for biological processes and enrichment analysis^78–80^. Data was extracted from the output dataset and graphed using R. Proteomic interactions of the same relative expression data were also classified through KEGG-based proteomapping software^81^ and are presented as obtained.

### miRNA Isolation

RNA was isolated from LVVs using the Total Exosome RNA & Protein Isolation Kit (Thermo Fisher Scientific). Briefly, isolated LVVs were resuspended in exosome resuspension buffer and incubated with an equal volume of denaturation solution at 4 °C for 5 min. The solution was then mixed with an equal volume of Acid-Phenol:Chloroform by vortexing for 30 seconds and centrifuged for 5 min at 15,000g. The resulting aqueous phase was extracted and combined with 1.25x volume of 100% ethanol, then transferred to the provided spin column. The spin column was centrifuged at 10,000g for 15 seconds to bind and wash the RNA, then the RNA was eluted in the provided elution solution and quantified via a microvolume spectrophotometer (Nanodrop 2000, Thermo Fisher Scientific).

### Profiling of Total miRNA Population

Immediately following isolation, the eluted miRNA was concentrated using 3 kDa microcentrifuge spin filters (Amicon). Briefly, the 100 µL miRNA solution was worked up to 420 µL with RNAse-free water and placed into a filter, then centrifuged at 14,000g for 90 minutes. Next, the filter was inverted into a fresh collection tube, and centrifuged at 8,000g for 2 minutes. The resulting isolate is 20-25 µL of concentrated miRNA, which was quantified by a microvolume spectrophotometer. Concentrated miRNA was then prepared for miRNA profiling (NanoString) according to the manufacturer’s protocol. Briefly, the provided miRNA codeset was mixed with the provided hybridization buffer to produce a master mix, and spike-in miRNA controls were prepared at 200 pm. In order, the master mix, concentrated sample miRNA, spike-in miRNA, and provided probes were mixed in a PCR plate and incubated at 65 °C for 16 h. The hybridized solution was then mixed with 15 µL of provided hybridization buffer, for a total volume of 30-35 µL, and added to the provided microfluidic cartridge. The assay was run with the provided protocol for total miRNA analysis, and data was processed and analyzed using the provided software using the recommended settings.

### miRNA PCR

Isolated miRNA content was quantified by real time quantitative PCR (RT-qPCR) using the miScript PCR Kit (Qiagen) with a CFX Connect Real-Time system (Bio-Rad). cDNA was prepared from 100 ng of RNA template, and the primers used were hsa-miR-125b-5p, hsa-miR-143-3p, hsa-miR-145-5p, hsa-miR-199a-3p, hsa-miR-221-3p, and hsa-miR-222-3p (Supplemental Table S7). Results were quantified relative to RNU6B, the recommended control. Non template controls were also used, and no signal was detected from these controls.

### miRNA Transfection

hCFs were seeded and wounded as above for the wound healing assay. After wounding, cells were incubated with DMEM miRNA transfection media with either a cocktail of mimics of identified miRNA of interest (miR-125, *ACGGGUUAGGCUCUUGGGAGCU*; miR-143, *UGAGAUGAAGCACUGUAGCUC*; miR-199a, *ACAGUAGUCUGCACAUUGGUUA*; miR-222, *AGCUACAUCUGGCUACUGGGU*; all miRVana) or a scramble miRNA control (Negative Control #2, Ambion) at 40 nM. Transfection media was prepared with Lipofectamine 3000 according to the manufacturer’s specifications. Imaging and analysis were performed as above. After wounding, the transfected cells were fixed, stained and imaged as above in immunostaining. Two independent experiments with N = 3 biological replicates each were performed, with the later fixed at 48 h due to the high rates of cell death observed beyond that time point.

### Hypoxia Assay

hCFs were seeded as above for miRNA transfection assays and transfected with the mentioned miRNAs for 24 hours to maximize miRNA uptake and minimize cell death. Randomly selected plates were also incubated with BrdU (Abcam, 10 µM) for proliferation assessment. Following this, hCFs were transferred to deoxygenated glucose-free media (RPMI, Thermo Fisher) and subjected to hypoxia for 3 hours, which we have previously shown is sufficient to induce MI-like cell death^77^. BrdU-treated plates were subsequently fixed and prepared for staining as described in immunostaining, and stained with anti-BrdU (Abcam), while non-BrdU plates were stained with calcein AM and ethidium homodimer-1 Live/Dead stains (Invitrogen, 1:1000) to quantify living and dead cells, respectively. Stained cells were imaged as in immunostaining, and images were quantified in ImageJ.

### miRNA Pathway Analysis

Nanostring data was normalized via Log10 normalization, and uploaded to the Clarivate MetaCore system for pathway analysis. miRNAs were identified by miRBase IDs. Analysis was conducted on the identified target miRNAs to construct a custom network. Automated network analysis was conducted with 50 nodes per network. Results were presented as pathways obtained from the software.

### Statistical Analysis

Results were analyzed by one-way analysis of variance (ANOVA) with post-hoc Tukey’s HSD, two-way ANOVA with post-hoc Tukey’s multiple comparison test, or a two-tailed Student’s t-test with Welch’s correction for unequal standard deviation. Values are presented as the mean ± standard deviation (SD) unless otherwise indicated, and differences were considered significant when p ≤ 0.05.

## ACKNOWLEDGEMENTS

The lyophilization of decellularized ECM was conducted at the Center for Environmental Science and Technology (CEST) at the University of Notre Dame. We thank the Biophysics Instrumentation (BIC) Core Facility for the use of Optima MAX-XP Tabletop Ultracentrifuge. The authors acknowledge the use of the Electron Microscopy Core of the Notre Dame Integrated Imaging Facility, a designated core of the NIH-funded Indiana Clinical and Translational Sciences Institute. The Nanoparticle Tracking Analysis was conducted using the NanoSight NS300 at the Harper Cancer Research Institute (HCRI) Tissue Core Facility. The schematics in some figures were created using BioRender.com We thank Stanley Cheng, Zorlutuna Lab manager, for assisting in proofreading this manuscript prior to submission

## AUTHOR CONTRIBUTIONS

G.R., G.B., J.Y., and P.Z. designed research, G.R. performed research, G.R. analyzed data, G.B. and P.Z. conducted review and editing, P.Z. provided funding, project administration, and resources, G.R. wrote the paper.

