## Supplemental Information for "Age and Sex-Dependent Differences in Human Cardiac Matrix-Bound Exosomes Modulate Fibrosis through Synergistic miRNA Effects"

**SUPPLEMENTAL FIGURES**

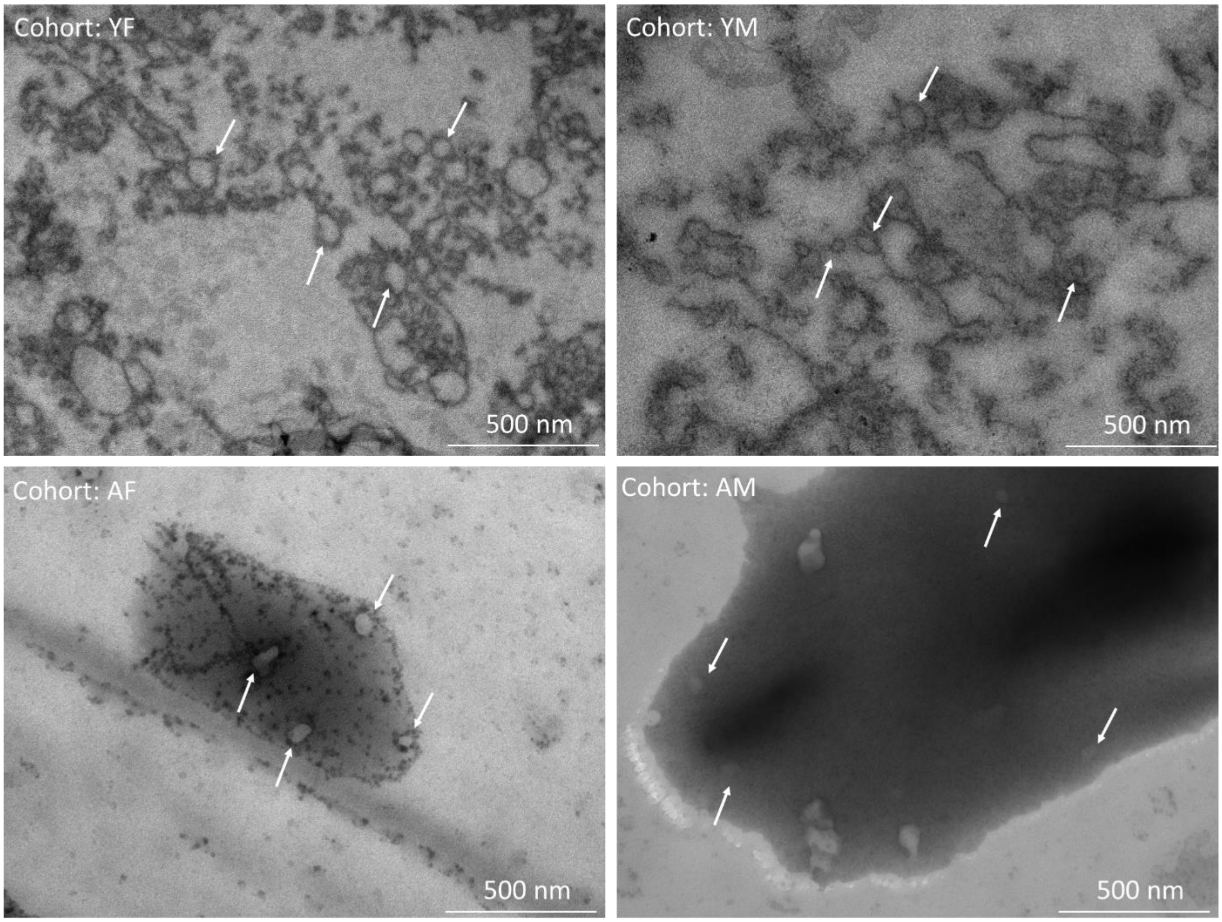


**Supplemental Figure S1**: LVVs Embedded in Extracellular Matrix. Representative images of LVVs embedded in ECM sections. ECM was sectioned and embedded in resin for imaging under TEM. While arrows indicate some, not all, LVVs in each image. Cohort is indicated in the top left of each panel.

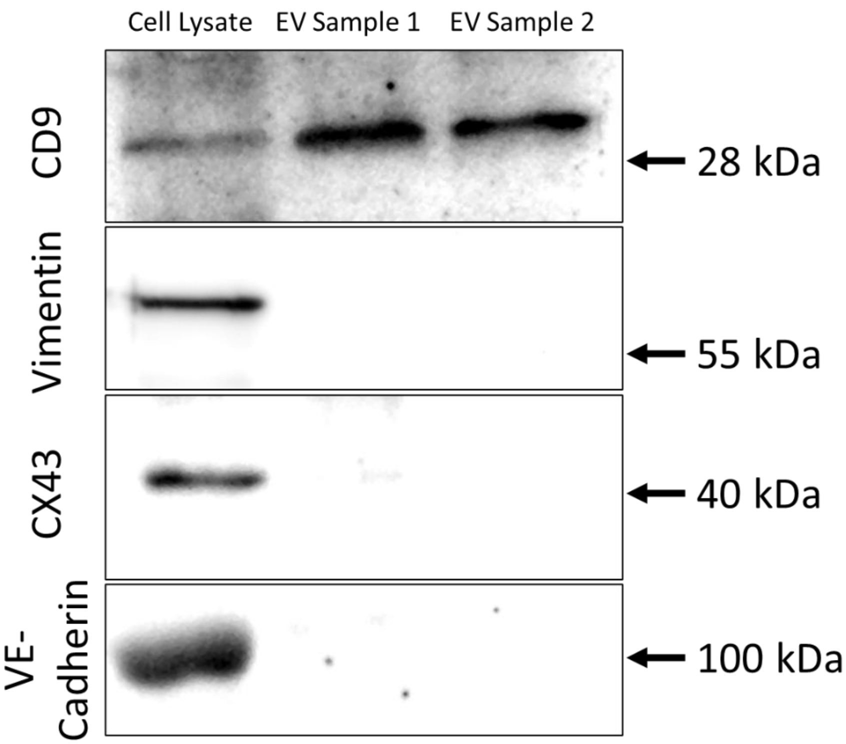


**Supplemental Figure S2**: LVVs Contain No Cell Debris Markers. Western blot images of CD9 and cell debris markers for cell lysate or 2 mixed EV samples. Cell lysate varied between marker, with CFs being used for CD9 and vimentin, cardiomyocytes for CX43, and endothelial cells for VE-cadherin. CD9 was expressed in the CF lysate, but no cell markers were shown in either EV blot.

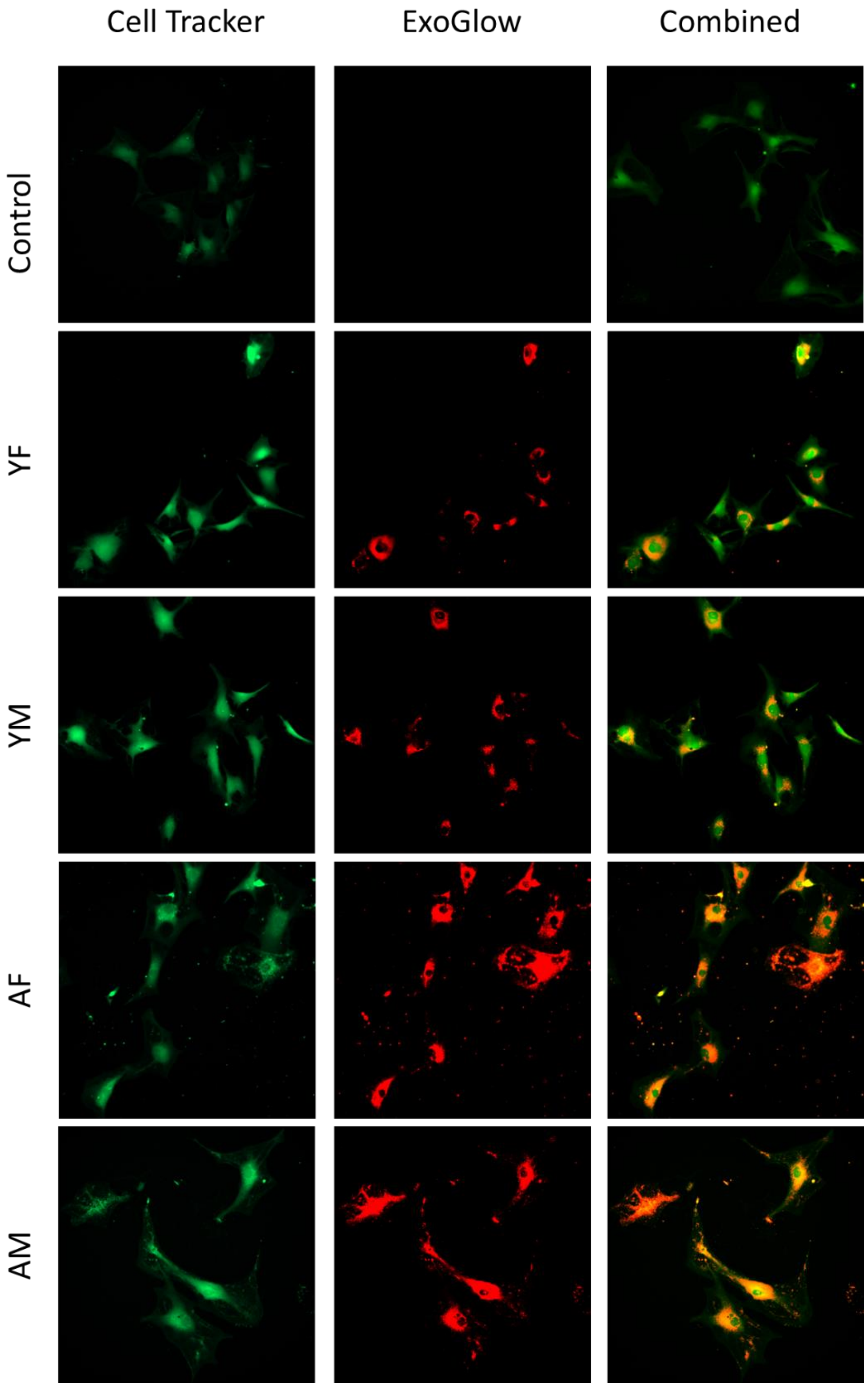


**Supplemental Figure S3**: Representative Images for EV Uptake Assay. Images for each cohort in the EV uptake assay at t = 24 h.

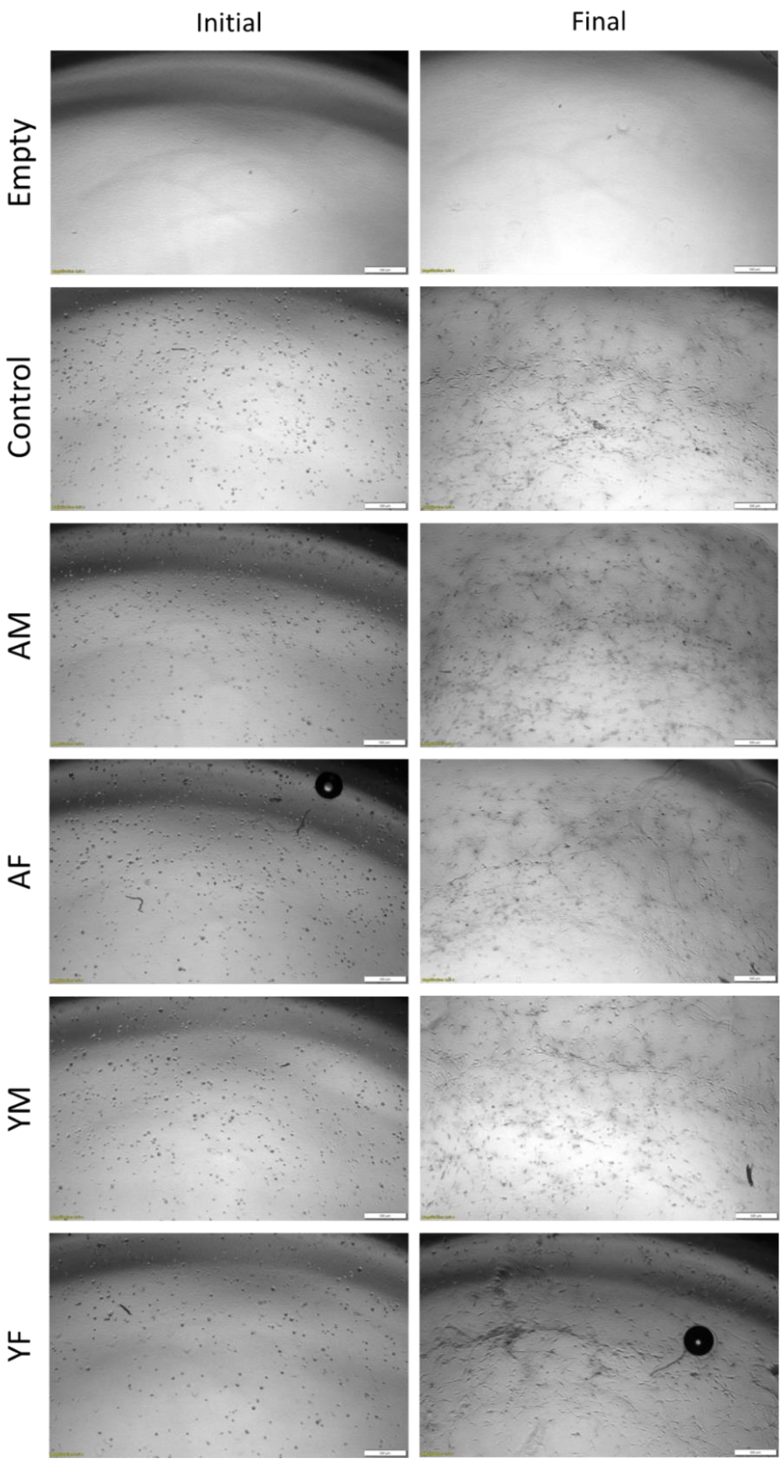


**Supplemental Figure S4**: Representative Images of hCFs in Collagen Gels for Seeding Density and Cell Attachment Visualization. Images at initial (t = 0 h) and final (t = 48 h) timepoints.

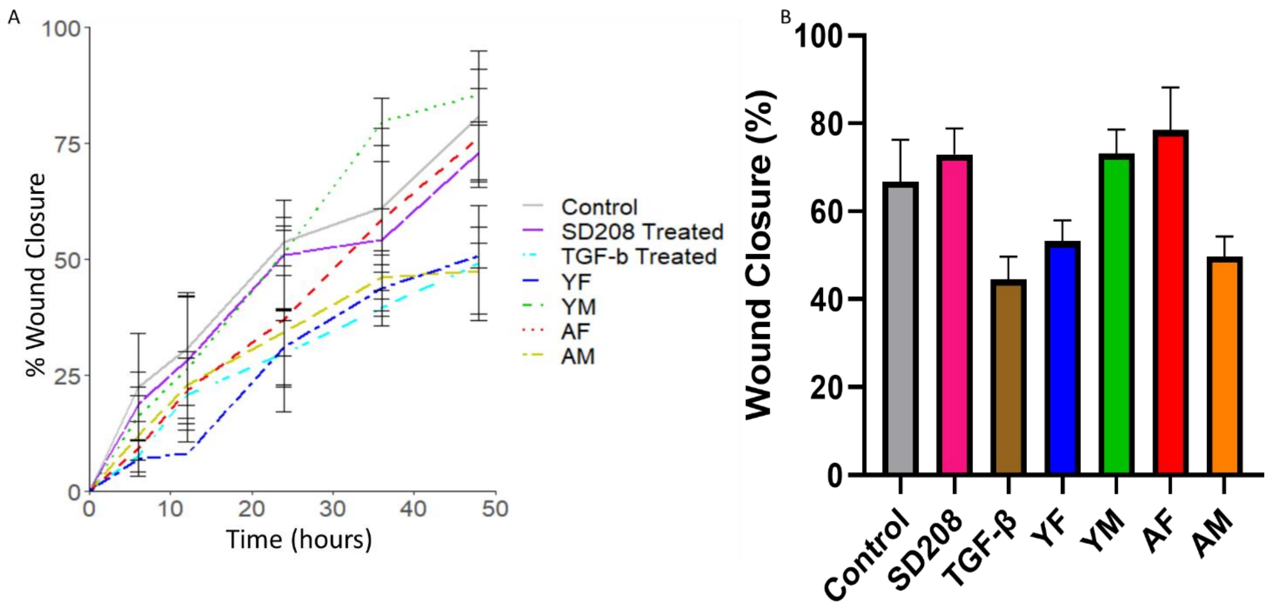


**Supplemental Figure S5**: Rat Cardiac Fibroblast Wound Healing Trial. Rate of wound closure percentage over 48 h (A) and comparison of wound closure at t = 48 h (B).

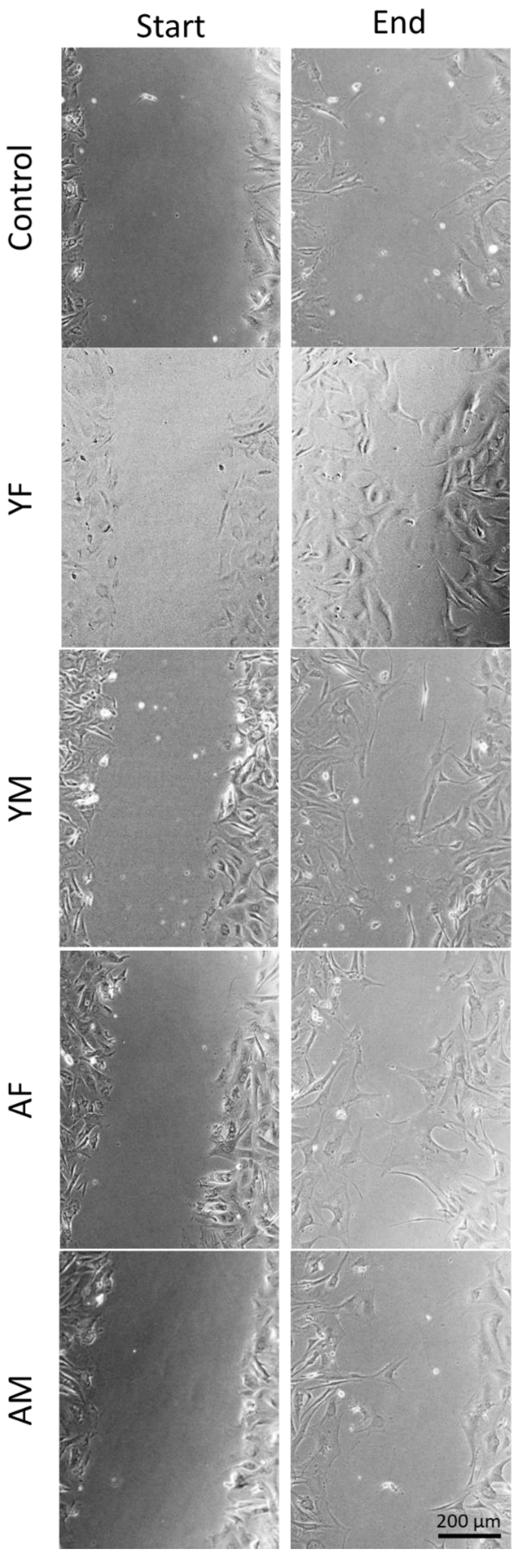


**Supplemental Figure S6**: Representative Images of hCF Wounds in the Wound Healing Assay. Images at initial (t = 0 h) and final (t = 96 h) timepoints.

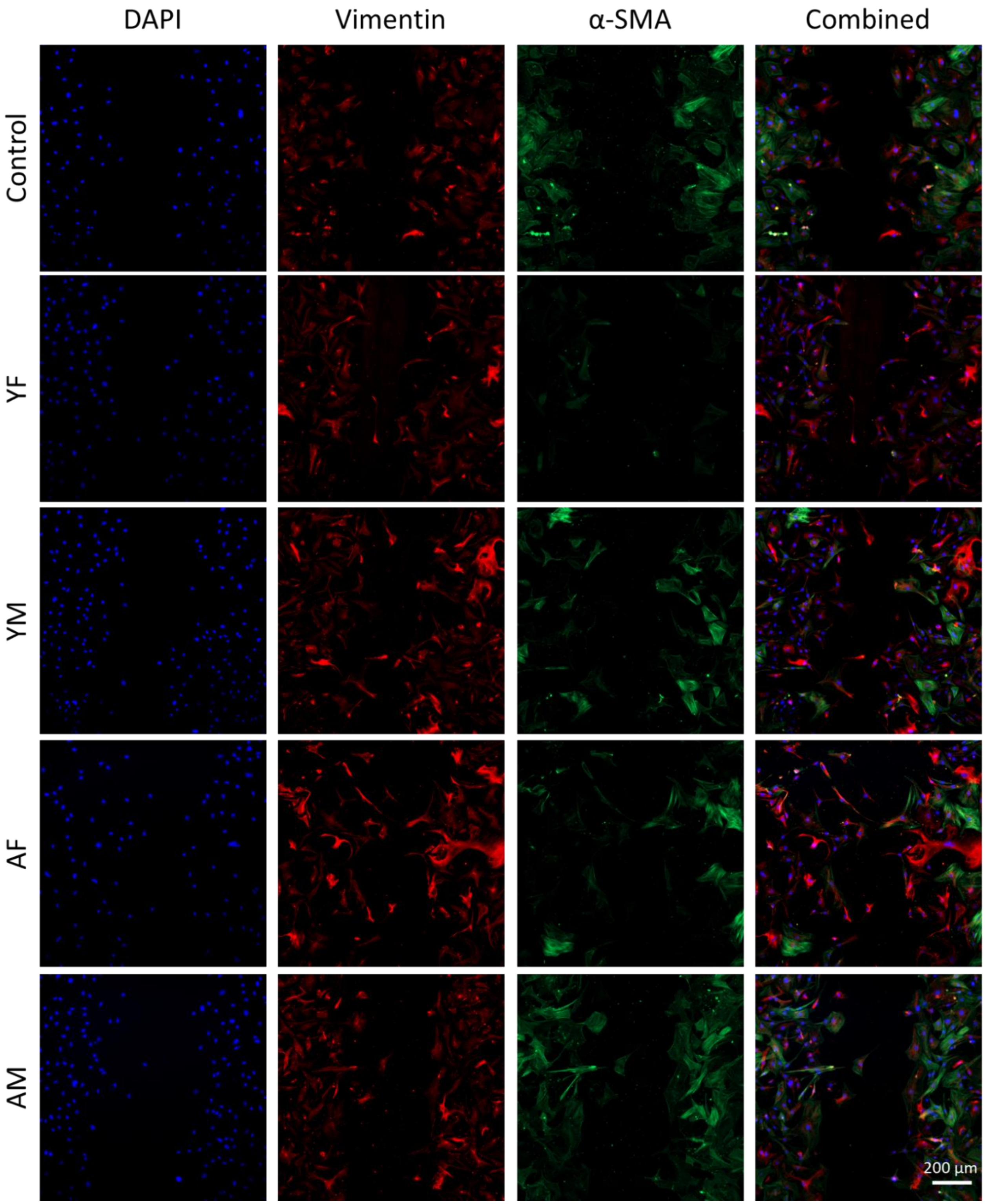


**Supplemental Figure S7**: Representative Images of hCF Wounds after Immunostaining. Images show DAPI (blue), Vimentin (red), $\alpha$-SMA (green), and combined channels.

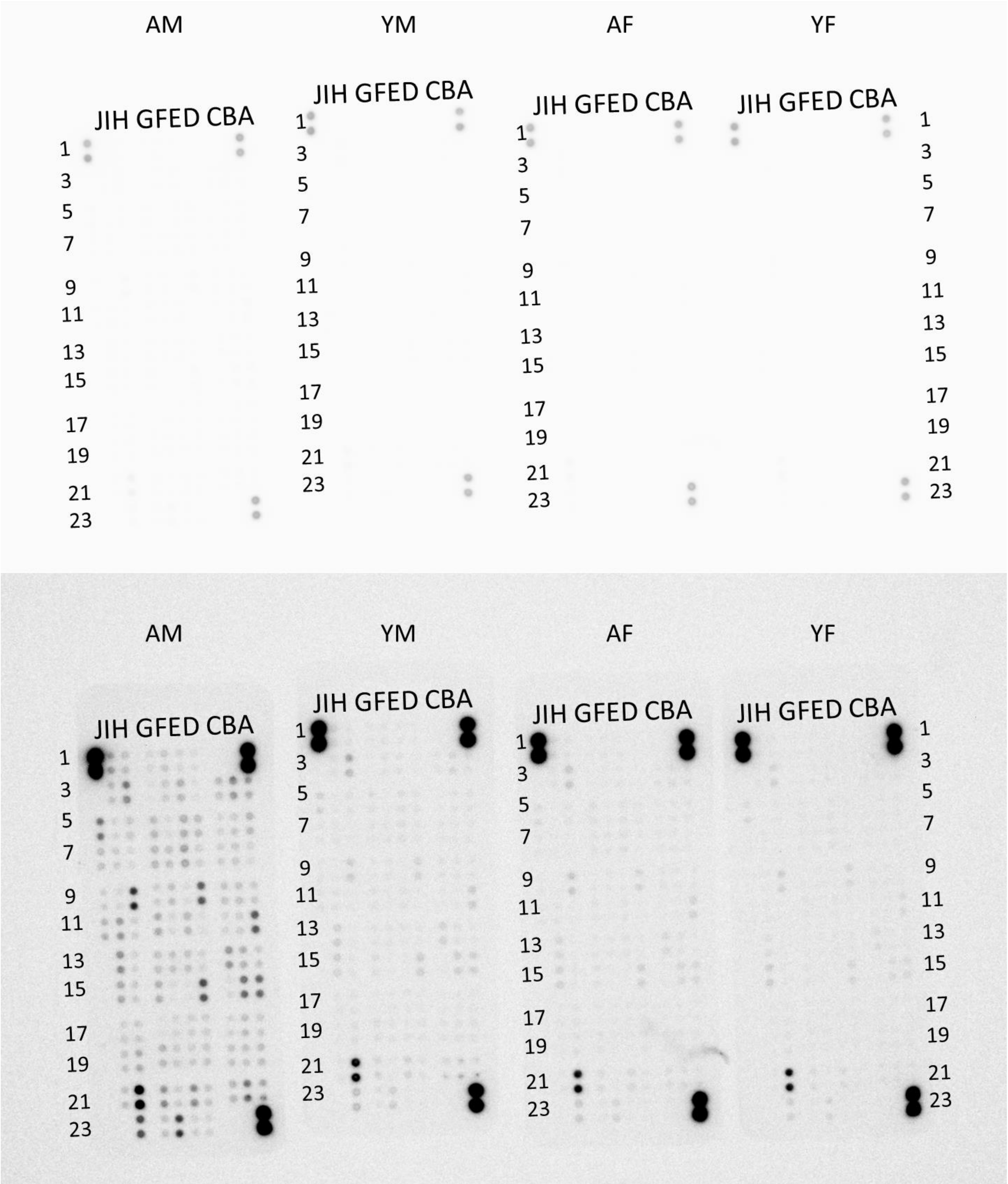


**Supplemental Figure S8**: Raw (top) and Adjusted (bottom) Blot Images from Human XL Cytokine Array.

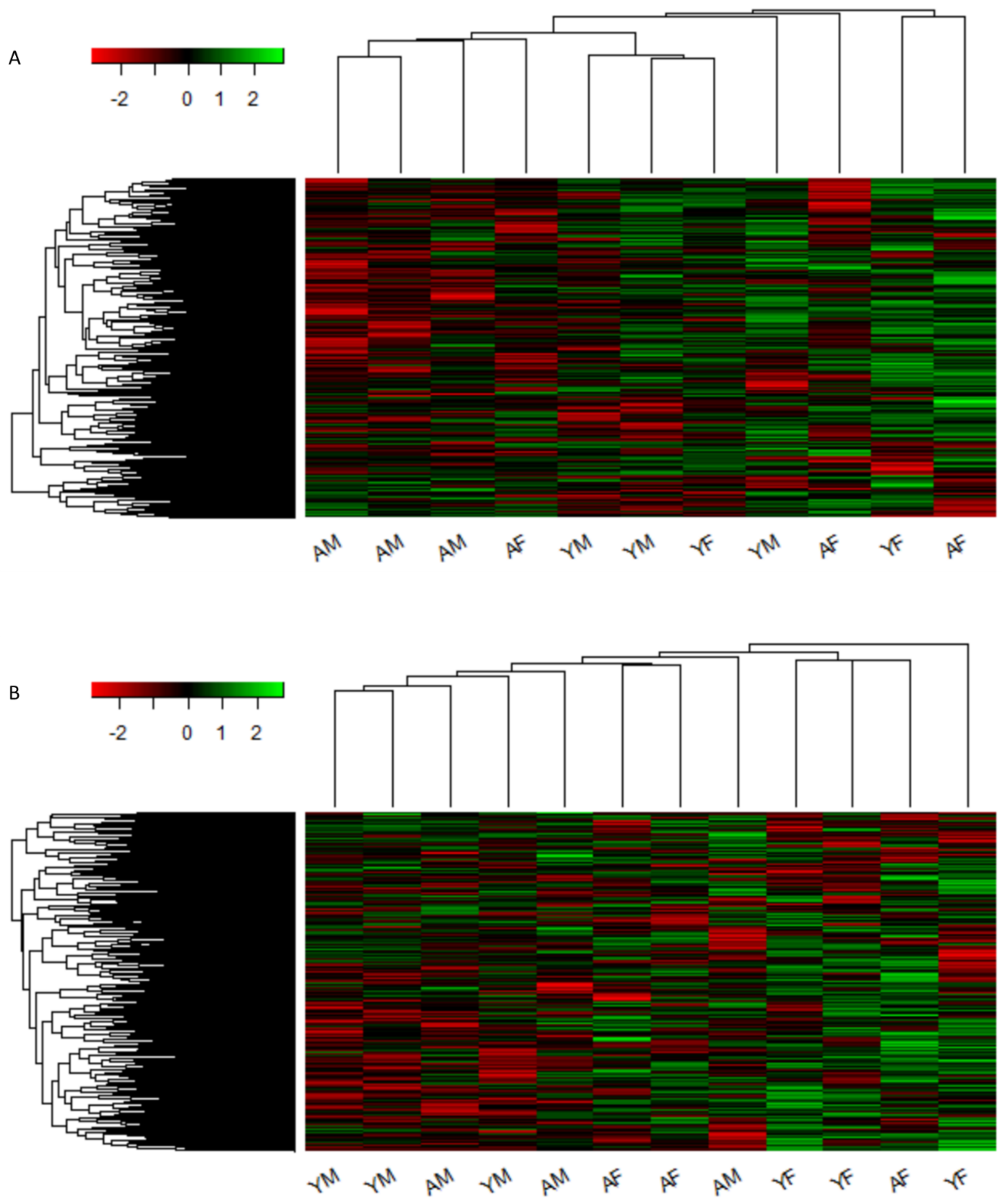


**Supplemental Figure S9**: Individual Replicate Heatmaps from Nanostring Analysis. Images show miRNA profile for individual replicates of human (A) or mouse (B) EVs. Clustering was performed in R using Euclidian distance calculations.

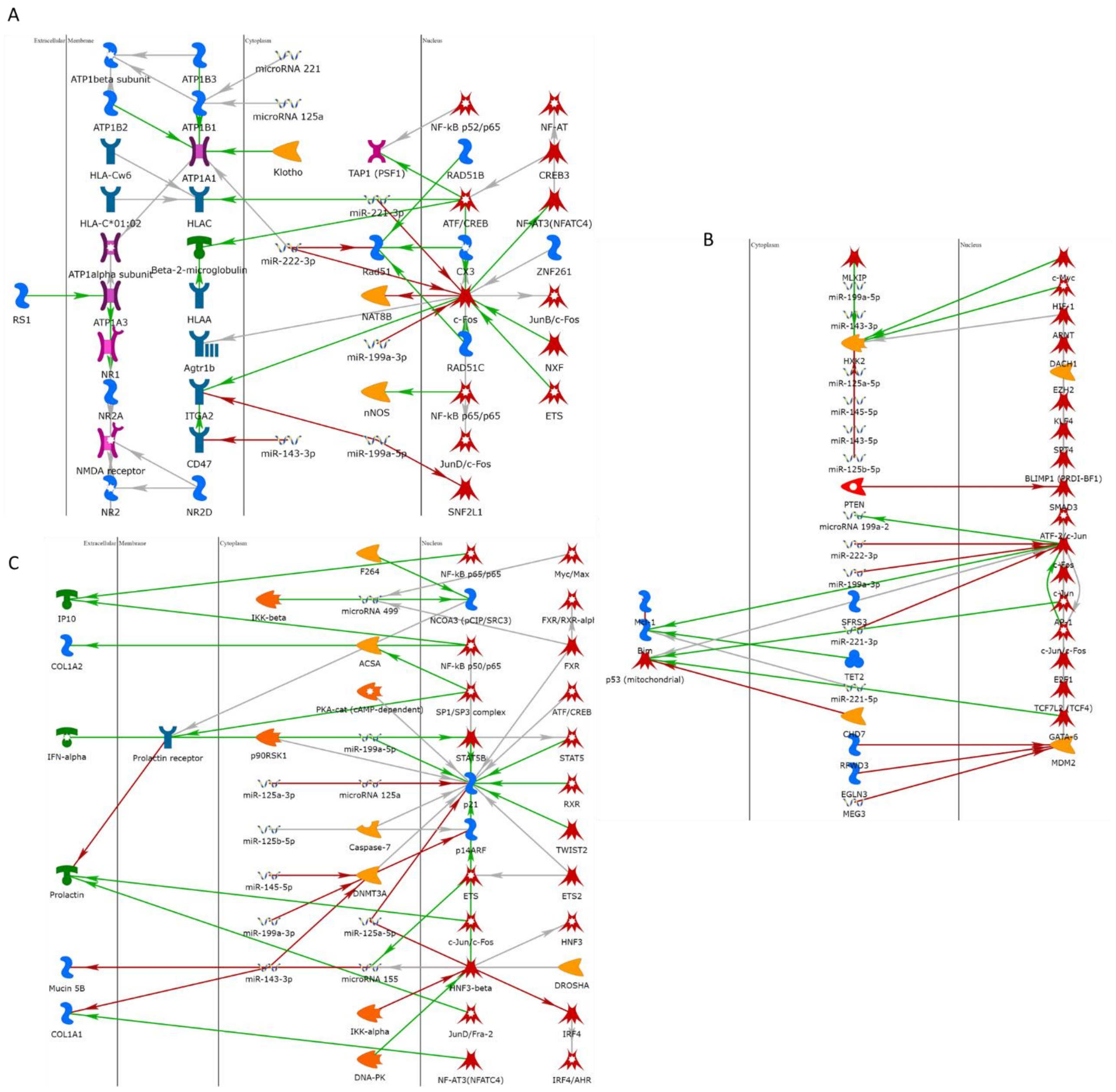

**Supplemental Figure S10**: Full Pathways from MetaCore Analysis. The full pathways provided from the MetaCore analysis performed on the miRNA data from human LVVs.

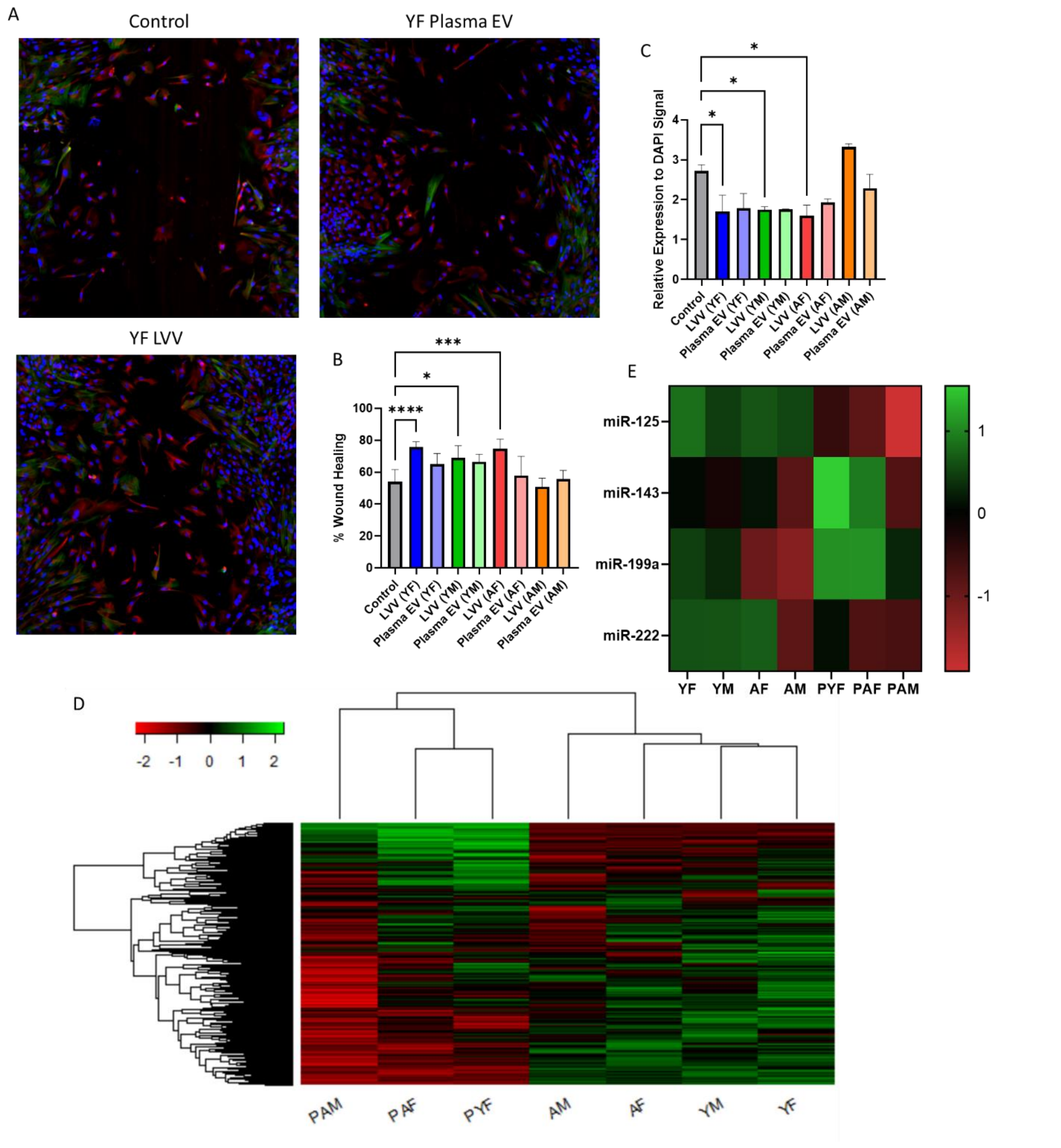


**Supplemental Figure S11**: Comparison of LVVs to plasma EVs. Comparison of results from iPSC-derived CFs for treatment with LVVs or plasma EVs. Performed assays were wound healing (A, B) and a-SMA immunostaining (A, C). Data are displayed with LVV and plasma shown side by side for each cohort. Also compared were Nanostring results (D) and relative expression of the identified target miRNAs (E). PYF = plasma YF, PAF = plasma AF, PAM = plasma AM.

### SUPPLEMENTAL TABLES

**Supplemental Table S1**: Cytokine Expression Table, Relative to YF.

| Protein | YM/YF | AF/YF | AM/YF | Protein | YM/YF | AF/YF | AM/YF | Protein | YM/YF | AF/YF | AM/YF |
|---|---|---|---|---|---|---|---|---|---|---|---|
| Adiponectin | 0.85 | 0.923 | 1.579 | IFN-γ | 0.912 | 0.96 | 1.169 | MCP1 | 0.982 | 1.006 | 1.421 |
| Apolipoprotein A-I | 0.957 | 1.023 | 1.678 | IGFBP2 | 0.918 | 0.98 | 1.276 | MCP3 | 0.992 | 1.023 | 1.904 |
| Angiogenin | 0.943 | 0.954 | 1.231 | IGFBP3 | 1.069 | 1.066 | 3.048 | M-CSF | 1.005 | 1.025 | 1.63 |
| Angiopoietin-1 | 0.962 | 0.976 | 1.383 | IL1α | 0.915 | 1.075 | 1.347 | MIF | 1.029 | 1.028 | 1.792 |
| Angiopoietin-2 | 1.149 | 1.117 | 2.994 | IL1β | 0.987 | 1.124 | 1.466 | MIG | 0.93 | 1.017 | 1.16 |
| BAFF | 0.927 | 0.975 | 1.389 | IL1ra | 0.957 | 1.015 | 1.395 | MIP1α/MIP1β | 1.034 | 1.044 | 1.719 |
| BDNF | 1.038 | 1.01 | 2.565 | IL2 | 0.893 | 1.002 | 1.248 | MIP3α | 1.032 | 1.019 | 2.082 |
| Complement component C5/C5a | 0.954 | 0.999 | 1.369 | IL3 | 0.947 | 1.008 | 1.281 | MIP3β | 0.984 | 1.021 | 1.539 |
| CD14 | 0.961 | 0.995 | 1.475 | IL4 | 0.925 | 0.97 | 1.771 | MMP9 | 0.972 | 1.033 | 1.62 |
| CD30 | 1.252 | 1.125 | 1.914 | IL5 | 1.012 | 1.008 | 1.87 | Myeloperoxidase | 1.403 | 1.201 | 2.18 |
| Reference Spots | 0.982 | 1.008 | 2.059 | IL6 | 1.008 | 1.026 | 2.034 | Osteopontin (OPN) | 1.033 | 1.056 | 1.526 |
| CD40 ligand | 0.911 | 0.956 | 1.31 | IL8 | 0.928 | 0.966 | 1.19 | PDGF-AA | 0.998 | 1.012 | 1.248 |
| Chitinase 3-like 1 | 0.915 | 0.991 | 1.409 | IL10 | 0.99 | 1.035 | 1.734 | PDGF-AB/BB | 1.034 | 1.156 | 3.213 |
| Complement factor D | 0.942 | 0.983 | 1.495 | IL11 | 0.962 | 1.055 | 1.559 | Pentraxin 3 | 0.996 | 1.03 | 1.43 |
| C-reactive protein | 0.898 | 0.965 | 1.309 | IL12 p70 | 0.956 | 1 | 1.175 | PF4 | 0.985 | 1.02 | 1.342 |
| Crypto-1 | 0.984 | 1.025 | 1.717 | IL13 | 0.979 | 1.036 | 1.4 | RAGE | 1 | 1.04 | 1.426 |
| Cystatin C | 1.09 | 1.051 | 2.581 | IL15 | 0.961 | 1.005 | 1.375 | RANTES | 0.989 | 1.042 | 1.398 |
| DKK-1 | 0.919 | 0.938 | 1.518 | IL16 | 1.023 | 1.061 | 2.084 | RBP4 | 1.004 | 1.027 | 1.772 |
| DPPIV | 0.933 | 1.028 | 1.346 | IL17A | 0.892 | 0.957 | 1.257 | Relaxin 2 | 1.154 | 1.116 | 1.411 |
| EGF | 1.155 | 1.051 | 2.455 | IL18Bpa | 0.996 | 1.015 | 1.492 | Resistin | 1.243 | 1.121 | 2.936 |
| EMMPRIN | 0.86 | 0.948 | 1.465 | IL19 | 1.03 | 1.008 | 1.499 | SDF1α | 1.107 | 1.166 | 2.149 |
| ENA-78 | 0.876 | 0.978 | 1.222 | IL22 | 0.949 | 1.026 | 1.252 | SERPINE1 | 1.054 | 1.033 | 1.634 |
| Endolgin | 0.936 | 0.99 | 1.291 | IL23 | 0.994 | 1.025 | 1.608 | SHBG | 1.022 | 1.026 | 1.365 |
| FAS ligand | 0.965 | 1.02 | 1.461 | IL24 | 0.987 | 1.005 | 1.55 | ST2 | 1.054 | 1.034 | 1.454 |
| FGFb | 0.881 | 0.957 | 1.224 | IL27 | 0.983 | 1.005 | 1.284 | TARC | 0.952 | 0.965 | 1.287 |
| FGF7 | 1.066 | 1.036 | 1.948 | IL31 | 0.996 | 1.039 | 1.555 | TFF3 | 1.028 | 1.07 | 2.245 |
| FGF19 | 0.886 | 0.982 | 1.199 | IL32 | 0.959 | 0.985 | 1.208 | TfR | 1.163 | 1.123 | 2.229 |
| FLT3 ligand | 0.886 | 0.956 | 1.186 | IL33 | 0.955 | 1.014 | 1.302 | TGFα | 0.927 | 0.961 | 1.451 |
| G-CSF | 0.907 | 1.097 | 1.36 | IL34 | 0.981 | 1.007 | 1.462 | Thrombospondin 1 | 1.038 | 1.066 | 1.448 |
| GDF15 | 1.057 | 1.042 | 1.556 | IP10 | 0.947 | 0.972 | 1.392 | TNFα | 0.982 | 1.031 | 1.559 |
| GM-CSF | 0.944 | 0.985 | 1.196 | I-TAC | 1.074 | 1.067 | 3.311 | uPAR | 0.997 | 1.039 | 1.582 |
| GROα | 0.919 | 0.962 | 1.121 | Kallikrein 3 | 0.996 | 0.994 | 1.332 | VEGF | 1.08 | 1.019 | 2.539 |
| Growth Hormone | 0.969 | 0.995 | 1.634 | Leptin | 0.946 | 0.998 | 1.186 | CD31 | 1.033 | 1.082 | 1.717 |
| HGF | 0.969 | 0.951 | 1.338 | LIF | 1.021 | 1.049 | 1.715 | TIM3 | 1.035 | 1.067 | 1.413 |
| ICAM-1 | 0.984 | 1.016 | 2.843 | Lipocalin 2 | 0.991 | 1.026 | 1.75 | VCAM1 | 1.085 | 1.052 | 1.832 |

**Supplemental Table S2**: PANTHER Gene Ontology Pathway Enrichment. ND = No detection.

| Pathway | ID | YF | YM | AF | AM |
|---|---|---|---|---|---|
| T-helper 17 cell lineage commitment | 0072540 | 3.706 | 3.304 | 3.373 | 0.000 |
| icosanoid metabolic process | 0032966 | 3.307 | 2.907 | 2.975 | 0.000 |
| alpha-beta T cell lineage commitment | 0002363 | 3.138 | 2.735 | 2.804 | 0.000 |
| negative regulation of blood pressure | 0045776 | 3.592 | 3.000 | 3.101 | 0.000 |
| renal protein absorption | 0071801 | 3.134 | 4.393 | 4.496 | 2.577 |
| collagen metabolic process | 0010713 | 3.246 | 4.554 | 4.658 | 2.686 |
| negative regulation of inflammatory response | 0050728 | 2.971 | 3.597 | 3.815 | 3.110 |
| programmed necrotic cell death | 0097300 | ND | 3.668 | 3.770 | 3.431 |
| positive regulation of ROS biosynthetic process | 1903428 | 0.000 | 5.470 | 4.244 | 5.076 |
| negative regulation of wound healing | 0061045 | ND | 3.644 | ND | 5.738 |
| positive regulation of receptor-mediated endocytosis | 0048260 | 3.408 | 4.080 | 4.216 | 6.388 |
| positive regulation of interleukin-1 production | 0032732 | 3.267 | 5.204 | 4.031 | 8.903 |
| positive regulation of interleukin-6 production | 0032755 | 5.521 | 8.469 | 5.979 | 9.127 |
| tissue remodeling | 0034103 | 3.058 | 6.179 | 5.032 | 9.767 |
| regulation of epithelial cell migration | 0010632 | 0.000 | 5.466 | 4.706 | 11.240 |
| monocyte chemotaxis | 0002548 | 7.06 | 7.590 | 9.452 | 11.845 |
| cellular response to growth factor stimulus | 0071356 | 8.080 | 9.597 | 11.25 | 15.717 |
| negative regulation of cell death | 0060548 | 5.400 | 9.757 | 9.597 | 17.735 |
| positive regulation of TNF production | 0032760 | 7.128 | 7.252 | 6.117 | 18.384 |
| regulation of TNF production | 0032680 | 7.189 | 6.967 | 6.019 | 19.604 |
| myeloid leukocyte migration | 0097529 | 8.959 | 14.186 | 13.16 | 23.492 |
| positive regulation of cytokine production | 0001819 | 10.378 | 11.818 | 10.223 | 24.118 |
| necroptotic signaling pathway | 0097527 | ND | ND | ND | 3.048 |
| positive regulation of T cell apoptotic process | 1901857 | ND | ND | ND | 3.252 |
| extracellular matrix disassembly | 0010716 | ND | ND | ND | 3.520 |
| chronic inflammatory response | 0002544 | ND | ND | ND | 4.231 |
| positive regulation of tissue remodeling | 0034105 | ND | ND | ND | 4.499 |
| regulation of chronic inflammatory response | 0002676 | ND | ND | ND | 4.602 |
| aging | 0007568 | ND | ND | ND | 5.140 |
| response to hypoxia | 0001666 | ND | ND | ND | 7.128 |

**Supplemental Table S3**: Ages of Mice Obtained for Tissue Collection, and Corresponding Human Age.

| Cohort | Mouse Age | Corresponding Human Age | Human Sample Age Range |
|---|---|---|---|
| **YF** | 16 weeks | 26 | 19-29 |
| **YM** | 16 weeks | 26 | 19-29 |
| **AF** | 72 weeks | 60-65 | 50-65 |
| **AM** | 72 weeks | 60-65 | 50-65 |

**Supplemental Table S4**: Additional miRNA Targets Identified from miRNA Profiling and Relative Expression Levels.

| Micro RNA | YF/AM | YM/AM | AF/AM |
|---|---|---|---|
| **miR-1306-5p** | 1.31 | 7.43 | 1.4 |
| **miR-374c-5p** | 1.63 | 5.01 | 4.7 |
| **miR-764** | 2.23 | 4.77 | 2.2 |
| **miR-520f-3p** | 3.46 | 4.68 | 2.61 |
| **miR-493-3p** | 5.67 | 4.34 | 4.7 |
| **miR-302e** | 3.63 | 3.79 | 4.47 |
| **miR-181a-5p** | 3.61 | 2.68 | 3.16 |
| **miR-3615** | 3.49 | 2.5 | 2.13 |
| **miR-18b-5p** | 3.45 | 2.24 | 1.4 |
| **miR-92a-3p** | 2.58 | 3.96 | 4.8 |
| **miR-660-5p** | 1.72 | 3.82 | 4.27 |
| **miR-606** | 1.2 | 3.31 | 3.69 |
| **miR-629-5p** | 2.18 | 1.77 | 3.69 |
| **miR-211-3p** | 1.75 | 3.07 | 3.32 |
| **miR-378i** | -2.02 | -2.99 | -1.69 |
| **miR-3605-5p** | -1.09 | -2.53 | 1.24 |
| **miR-324-5p** | -1.57 | -2.4 | -1.81 |
| **miR-208b-3p** | -1.09 | -2.4 | 1.04 |
| **miR-1252-5p** | -1.34 | -2.29 | 1.19 |
| **miR-1270** | -1.52 | -2.29 | 1.09 |
| **miR-30a-3p** | 1.19 | -2.16 | -1.06 |
| **miR-1293** | 1.11 | -2.15 | -1.59 |
| **miR-30a-5p** | 1.22 | -2.11 | -1.01 |
| **miR-142-3p** | -1.11 | -2.1 | -2.37 |
| **miR-607** | -3.75 | -1.34 | 1.05 |
| **miR-892b** | -2.73 | -1.38 | -1.31 |
| **miR-1307-5p** | -2.56 | 1.23 | -1.06 |
| **miR-601** | -2.43 | 1.26 | -1.27 |
| **miR-651-3p** | -2.36 | -1.65 | 1.14 |
| **miR-296-5p** | -2.16 | 1.4 | 1.48 |
| **miR-92a-1-5p** | -2.14 | 1.04 | -2 |
| **miR-1244** | 1.44 | 1.62 | -4.69 |
| **miR-550a-5p** | -1.19 | 1.35 | -4.54 |
| **miR-508-3p** | 1.04 | 1.25 | -3.83 |
| **miR-4524a-5p** | 1.08 | 1.32 | -3.38 |

**Supplemental Table S5**: Age, Sex, Cardiac Involvement in Cause of Death, and Preconditions for Human Heart Donors.

| Cohort | Age (years) | Heart-related Cause of Death | CVD Precondition(s) |
|---|---|---|---|
| **YF** | 24 | No | N/A |
| **YF** | 29 | No | Obesity |
| **YM** | 19 | No | N/A |
| **YM** | 27 | No | N/A |
| **YM** | 40 | No | Hypertension |
| **AF** | 52 | No | Valve Stenosis |
| **AF** | 56 | No | N/A |
| **AF** | 60 | No | Hypertension |
| **AM** | 63 | No | Hypertension |
| **AM** | 66 | No | N/A |
| **AM** | 51 | No | N/A |

**Supplemental Table S5**: Human Cytokines Tested in Human XL Array Blot Assay and Corresponding Blot Location.

| Spot Location | Protein | Spot Location | Protein | Spot Location | Protein |
|---|---|---|---|---|---|
| **A1, A2** | Reference Spots | **D11, D12** | IGFBP2 | **G13, G14** | MIF |
| **A3, A4** | Adiponectin | **D13, D14** | IGFBP3 | **G15, G16** | MIG |
| **A5, A6** | Apolipoprotein A-I | **D15, D16** | IL1$\alpha$ | **G17, G18** | MIP1$\alpha$/MIP1$\beta$ |
| **A7, A8** | Angiogenin | **D17, D18** | IL1$\beta$ | **G19, G20** | MIP3$\alpha$ |
| **A9, A10** | Angiopoietin-1 | **D19, D20** | IL1ra | **G21, G22** | MIP3$\beta$ |
| **A11, A12** | Angiopoietin-2 | **D21, D22** | IL2 | **G23, G24** | MMP9 |
| **A13, A14** | BAFF | **D23, D24** | IL3 | **H1, H2** | Myeloperoxidase |
| **A15, A16** | BDNF | **E1, E2** | IL4 | **H3, H4** | Osteopontin (OPN) |
| **A17, A18** | Complement component C5/C5a | **E3, E4** | IL5 | **H5, H6** | PDGF-AA |
| **A19, A20** | CD14 | **E5, E6** | IL6 | **H7, H8** | PDGF-AB/BB |
| **A21, A22** | CD30 | **E7, E8** | IL8 | **H9, H10** | Pentraxin 3 |
| **A23, A24** | Reference Spots | **E9, E10** | IL10 | **H11, H12** | PF4 |
| **B3, B4** | CD40 ligand | **E11, E12** | IL11 | **H13, H14** | RAGE |
| **B5, B6** | Chitinase 3-like 1 | **E13, E14** | IL12 p70 | **H15, H16** | RANTES |
| **B7, B8** | Complement factor D | **E15, E16** | IL13 | **H17, H18** | RBP4 |
| **B9, B10** | C-reactive protein | **E17, E18** | IL15 | **H19, H20** | Relaxin 2 |
| **B11, B12** | Crypto-1 | **E19, E20** | IL16 | **H21, H22** | Resistin |
| **B13, B14** | Cystatin C | **E21, E22** | IL17A | **H23, H24** | SDF1$\alpha$ |
| **B15, B16** | DKK-1 | **E23, E24** | IL18Bpa | **I1, I2** | SERPINE1 |
| **B17, B18** | DPPIV | **F1, F2** | IL19 | **I3, I4** | SHBG |
| **B19, B20** | EGF | **F3, F4** | IL22 | **I5, I6** | ST2 |
| **B21, B22** | EMMPRIN | **F5, F6** | IL23 | **I7, I8** | TARC |
| **C3, C4** | ENA-78 | **F7, F8** | IL24 | **I9, I10** | TFF3 |
| **C5, C6** | Endolgin | **F9, F10** | IL27 | **I11, I12** | TfR |
| **C7, C8** | FAS ligand | **F11, F12** | IL31 | **I13, I14** | TGF$\alpha$ |
| **C9, C10** | FGFb | **F13, F14** | IL32 | **I15, I16** | Thrombospondin 1 |
| **C11, C12** | FGF7 | **F15, F16** | IL33 | **I17, I18** | TNF$\alpha$ |
| **C13, C14** | FGF19 | **F17, F18** | IL34 | **I19, I20** | uPAR |
| **C15, C16** | FLT3 ligand | **F19, F20** | IP10 | **I21, I22** | VEGF |
| **C17, C18** | G-CSF | **F21, F22** | I-TAC | **I23, I24** | Reference Spots |
| **C19, C20** | GDF15 | **F23, F24** | Kallikrein 3 | **J1, J2** | Reference Spots |
| **C21, C22** | GM-CSF | **G1, G2** | Leptin | **J3, J4** | Vitamin D BP |
| **D1, D2** | GRO$\alpha$ | **G3, G4** | LIF | **J5, J6** | CD31 |
| **D3, D4** | Growth Hormone | **G5, G6** | Lipocalin 2 | **J7, J8** | TIM3 |
| **D5, D6** | HGF | **G7, G8** | MCP1 | **J9, J10** | VCAM1 |
| **D7, D8** | ICAM-1 | **G9, G10** | MCP3 | **J23, J24** | Negative Control |
| **D9, D10** | IFN-$\gamma$ | **G11, G12** | M-CSF | | |

**Supplemental Table S6**: miRNA Primer Target Sequence and Target miRNA Sequence

| **Name** | ***Target***/Sequence |
|---|---|
| ***hsa-miR-125b-5p Primer*** | ***5’ACGGGUUAGGCUCUUGGGAGCU*** |
| hsa-miR-125b-5p | ACGGGUUAGGCUCUUGGGAGCU |
| mmu-miR-125b-5p | ACGGGUUAGGCUCUUGGGAGCU |
| ***hsa-miR-143-3p Primer*** | ***5’UGAGAUGAAGCACUGUAGCUC*** |
| hsa-miR-143-3p | UGAGAUGAAGCACUGUAGCUC |
| mmu-miR-143-3p | UGAGAUGAAGCACUGUAGCUC |
| ***hsa-miR-145-5p Primer*** | ***5’GUCCAGUUUUCCCAGGAAUCCCU*** |
| hsa-miR-145-5p | GUCCAGUUUUCCCAGGAAUCCCU |
| mmu-miR-145-5p | GUCCAGUUUUCCCAGGAAUCCCU |
| ***hsa-miR-199a-3p Primer*** | ***5’ACAGUAGUCUGCACAUUGGUUA*** |
| hsa-miR-199a-3p | ACAGUAGUCUGCACAUUGGUUA |
| mmu-miR-199a-3p | ACAGUAGUCUGCACAUUGGUUA |
| ***hsa-miR-221-3p Primer*** | ***5’AGCUACAUUGUCUGCUGGGUUUC*** |
| hsa-miR-221-3p | AGCUACAUUGUCUGCUGGGUUUC |
| mmu-miR-221-3p | AGCUACAUUGUCUGCUGGGUUUC |
| ***hsa-miR-222-3p Primer*** | ***5’AGCUACAUCUGGCUACUGGGU*** |
| hsa-miR-222-3p | AGCUACAUCUGGCUACUGGGU |
| ***mmu-miR-222-3p Primer*** | ***5’AGCUACAUCUGGCUACUGGGUCU*** |
| mmu-miR-222-3p | AGCUACAUCUGGCUACUGGGUCU |
